# Spatiotemporal Analysis of Remyelination Reveals a Concerted Interferon-Responsive Glial State That Coordinates Immune Infiltration

**DOI:** 10.1101/2025.04.22.649486

**Authors:** Michael-John Dolan, Raphael Rakosi-Schmidt, Eric Garcia, Yuesang Lin, Naeem M. Nadaf, Jessica Dixon, Shiran Guo, Nader Morshed, Constanze Depp, Jordan Doman, Judy Xia, Henna Jäntti, Beth Stevens, Evan Z. Macosko

**Affiliations:** The Stanley Center for Psychiatric Research, Broad Institute of MIT and Harvard, Cambridge, MA 02142, USA; F.M. Kirby Neurobiology Center, Boston Children’s Hospital, Boston, MA 02115, USA; Department of Neurobiology, Harvard Medical School, Boston, MA, USA; Society of Fellows, Harvard University, Cambridge, MA 02138, USA; Howard Hughes Medical Investigator, Boston Children’s Hospital, Boston, MA 02115, USA; Department of Psychiatry, Massachusetts General Hospital, Boston, MA, USA; Present address: Smurfit Institute of Genetics, Trinity College Dublin, Dublin 2, Ireland

## Abstract

Remyelination, the process by which axons are re-encased in myelin after injury, is a critical step in restoring brain function, yet the dynamics from initial injury to repair remain poorly characterized. Here, we combined optimized single-nucleus RNA-seq with Slide-seqv2, a high-resolution spatial transcriptomics technology, to densely reconstruct the cellular processes that coordinate remyelination after a focal demyelinating injury. This revealed several findings: First, we found extensive transcriptional diversity of glia and monocyte-derived macrophages from demyelination to repair. Second, we identified a population of infiltrating peripheral lymphocytes—predominantly CD8 T-cells and natural killer cells—that are enriched specifically during remyelination. Third, we identified a concerted interferon-response gene signature that is shared across several cell types—microglia, astrocytes, and the oligodendrocyte lineage—just prior to reestablishment of myelin. These interferon-responsive glia (IRG) form clusters around remyelinating white matter and their formation is solely dependent on the type I interferon receptor. Functionally, we found that IRG secrete the cytokine CXCL10 which mediates infiltration of peripheral lymphocytes into the repairing white matter. Depletion of the most abundant infiltrating lymphocyte, CD8 T-cells, attenuated the differentiation of mature oligodendrocytes during remyelination. Together, our data reveals the diversity of glial-immune interactions that orchestrate white matter repair and a type I-dependent glial state that drives lymphocyte influx into damaged white matter to modulate oligodendrocyte differentiation.

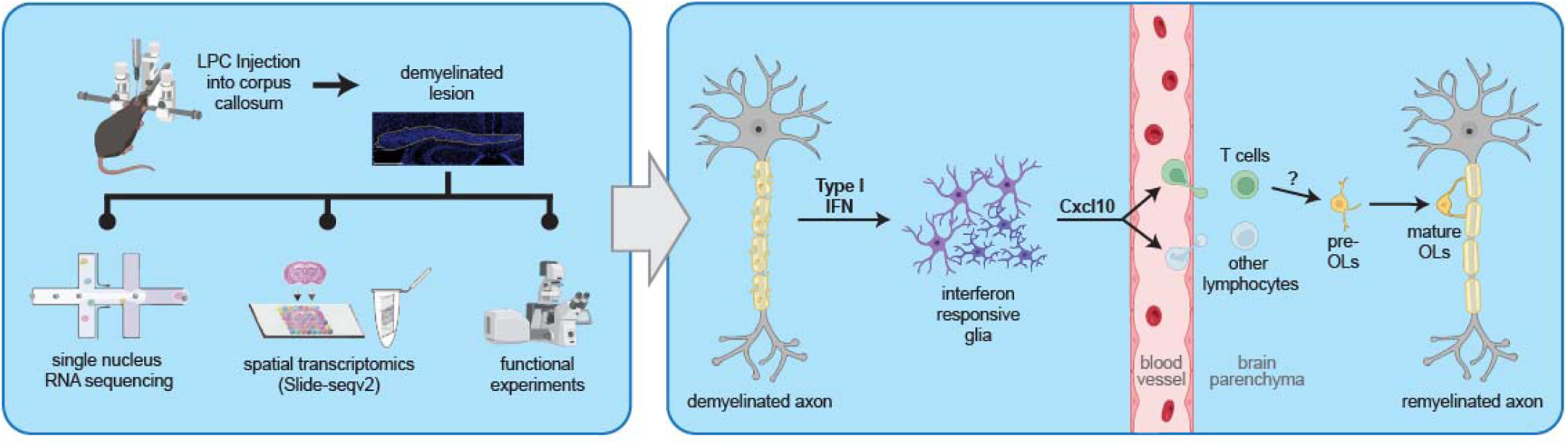

## Introduction

Myelin plays critical roles in neuronal communication, axonal health and metabolism^1–3^. Long-term loss of myelin results in dysfunctional neuronal activity, axonal damage and cell death^4^, making efficient maintenance and repair of myelin critical for brain health. Additionally, demyelination occurs in numerous neurodegenerative disorders such as Alzheimer’s disease^5,6^, multiple sclerosis^7^ and traumatic brain injury^8^, positioning remyelination as a target pathway to arrest or reverse neurodegeneration^1,5^. While axons are ultimately myelinated by oligodendrocytes, remyelination is profoundly dependent upon broader interactions between glial and immune cell types to clear debris and guide the differentiation of oligodendrocyte lineage cells at the site of damage^1–3,9–11^.

Single-cell RNA sequencing of the rodent and human central nervous system has revealed that glial cell types are heterogenous: microglia^12–14^, astrocytes^15^ and oligodendrocytes^16–18^ all adopt distinct transcriptional subtypes called “states”, especially in disease or after injury. Recent work has demonstrated that white matter damage results also in the formation of distinct glial subtypes^19,20^ and that these can play essential roles in tissue repair. For example, during remyelination distinct microglia signatures appear during damage and repair, with both playing essential roles in driving myelination^21–23^. Yet, the factors that drive the formation of these glial states, in addition to their downstream functions remain unknown, particularly in remyelination.

To resolve this, we combined single-nucleus and spatial transcriptomics to comprehensively profile the process of remyelination in the lysophosphatidylcholine (LPC) injection model, over a three-week period from injury to repair. Importantly, we optimized our nuclear extraction protocol to maximize the acquisition of rare immune cell types from small focal white matter lesions in the corpus callosum. This dataset revealed three major findings. First, we uncovered complex temporal transcriptional dynamics in all major glial classes throughout repair. Second, we identified sequential infiltration of leukocytes, initially monocyte-derived macrophages followed by CD8 T-and NK cells, during the remyelination phase. Third, we discovered a concerted interferon response gene signature in microglia, astrocytes, and oligodendrocytes. We found that these interferon-responsive glia (IRG) form clusters around repairing white matter and determined the formation of this state is solely dependent on the type I interferon receptor. To understand the function of the IRG state we perturbed an IRG-specific cytokine, CXCL10, and found that this promotes the parenchymal infiltration of peripheral lymphocytes. Moreover, we found that acutely depleting the most abundant of these infiltrating populations, CD8^+^ T-cells, attenuated oligodendrocyte maturation during remyelination. Together, our data reveal the extensive diversity of glial-immune interactions that orchestrate white matter repair and demonstrate how molecularly defined glial states and peripheral immune populations regulate oligodendrocyte differentiation during remyelination.

## Results

### A single-nucleus and spatial transcriptomic molecular census of remyelination

To model remyelination, we injected lysophosphatidylcholine (LPC), an endogenous phospholipid that causes demyelination^21^, into the corpus callosum. This results in acute myelin loss followed by remyelination in a stereotyped manner over 3 weeks, enabling dense reconstruction of this reparative process^21^. We chose to sample cellular states at four time points, representing distinct repair phases: initial myelin damage (3 days post injection, dpi), oligodendrocyte differentiation (7dpi), early (12 dpi) and late (18dpi) remyelination^21,24,25^ (Fig. 1a). To profile the gene expression of cells during remyelination, we developed an optimized extraction protocol to maximize the yield of nuclei from small white matter samples (Extended Data Fig. 1a) and performed single-nucleus RNA sequencing (snRNAseq)^26,27^ at all four time points with matched saline-injected controls (n=3 per time point and condition) (Fig. 1a). After filtration and quality control, we recovered 77,829 nuclear profiles consisting of oligodendrocytes, oligodendrocyte precursor cells (OPCs), astrocytes, microglia, endothelial cells, in addition to a small number of fibroblasts and mural cells (Fig. 1b-d, see Extended Data Fig. 1b for quality control metrics and Supplementary Table 1 for cell number per type). We also observed the infiltration of both monocyte-derived macrophages and lymphocytes exclusively in LPC-injected white matter (Fig. 1b-d).

**Figure 1:**
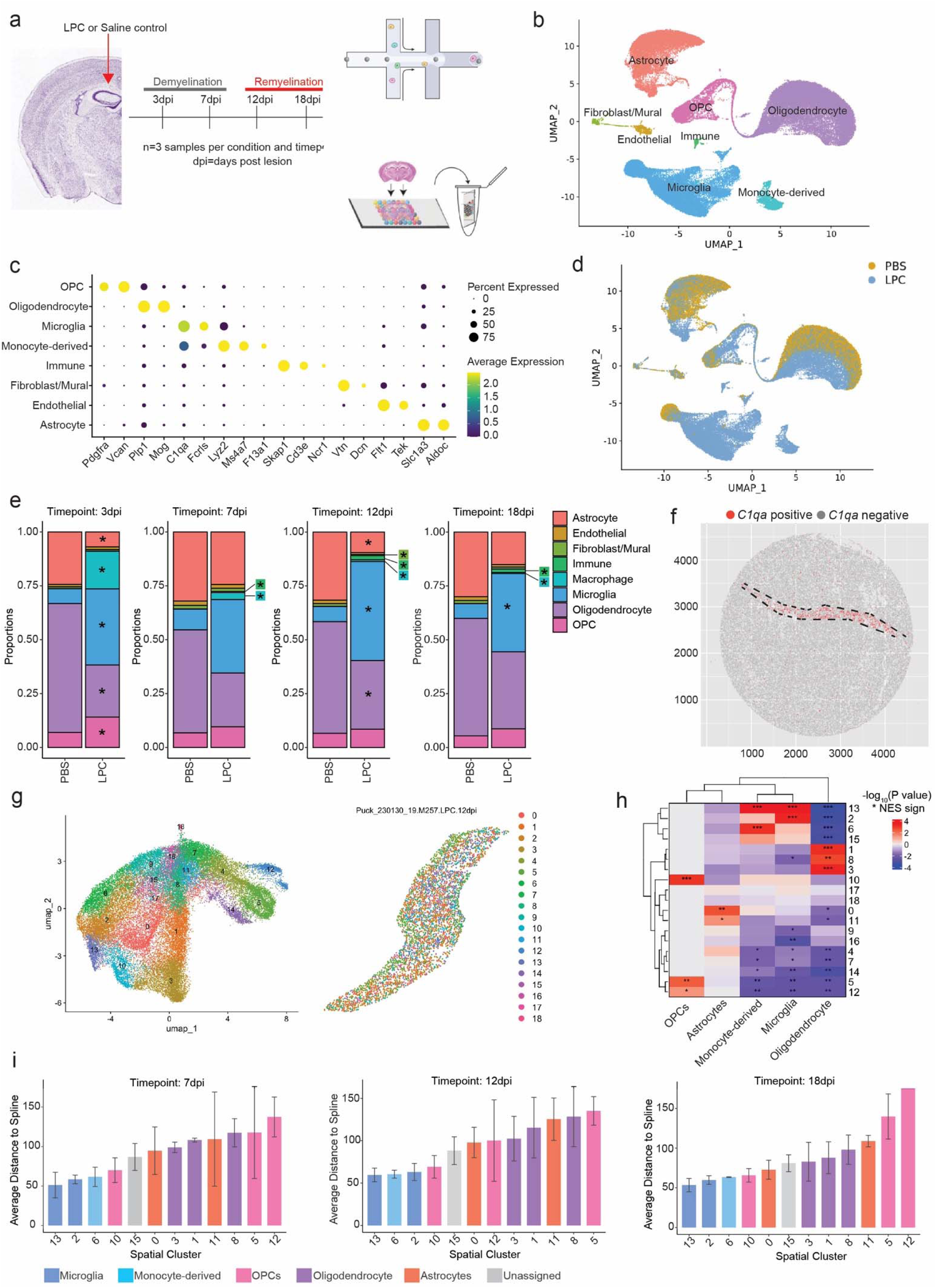
A single-cell and spatial transcriptomic atlas of remyelination. a) Schematic of the de-and remyelination dataset. PBS= Saline, LPC = lysophosphatidylcholine. b) Uniform Manifold Approximation and Projection (UMAP) projection of all glial cells; total of 77,829 nuclei, colored by cell type. c) Dot plot of cell type marker genes d) UMAP of all glial cells, colored by condition (PBS saline or LPC). e) Proportion of each cell type per time point and condition, * signifies statistical significance (p-value < 0.05) determined by a differential abundance test (Methods). f) Representative unsegmented Slide-seqv2 array with beads binarized by *C1qa* expression, revealing dense buildup of microglia in repairing white matter. Lesion outlined with a dashed line. g) (Left) UMAP projection of all bead transcriptomes across all Slide-seqv2 arrays. (Right) Representative spatial position of beads colored by cluster for 12dpi segmented lesion. h) Heatmap illustrating the relative enrichment significance (determined by gene set enrichment analysis) of positively enriched marker genes for each cell type (from the snRNAseq dataset) within all differentially expressed genes for each spatial cluster. * = p-value<0.05, ** = p-value<0.01 and *** = p-value<0.001. i) Average distance of beads of a particular spatial cluster from a spline through the lesion core. Plotted for 7dpi (left), 12dpi (middle) or 18dpi (right).

We first characterized the cellular composition of the lesion across time by calculating differential abundance at of each cell type in each of the four timepoints^28^ (Fig. 1e, Extended Data Fig. 1c and Supplementary Table 2). As expected, the proportion of oligodendrocytes was significantly decreased after demyelination compared to the saline control, and then gradually increased during each successive repair stage. This was contrasted with an extensive expansion of microglia at every assessed time point of de-and remyelination. The proportion of OPCs increased after initial demyelination, matching prior work showing this cell type undergoes proliferation and migration to the site of injury^25^.

To understand the spatial distribution of these cell types in white matter repair, we performed Slide-seqv2^30,31^, a transcriptome-wide, 10 micron resolution spatial assay, on 15 tissue sections taken throughout remyelination, matching the single-nucleus profiling above. We cropped only lesioned white matter (Fig. 1f) and identified 19 different spatial clusters (Fig. 1g, Extended Data Fig. 2a-c and Supplementary Table 3). To map cell type identities to these spatial clusters, we generated cell-type-specific gene lists using our snRNAseq data and measured the enrichment across clusters using Gene Set Enrichment Analysis (GSEA) or bead-based module scoring^32,33^ (Fig. 1h, Extended Data Fig. 2d and Supplementary Table 4). In both cases, we were able to map cell types to the same spatial clusters, including microglia (13,2,6), astrocytes (0, 11), OPCs (5,12, 10) and oligodendrocytes (1, 8, 3) (Fig. 1h). Clusters mapping to neurons were excluded because they mapped to the adjacent grey matter (Extended Data Fig. 1g). To understand how these clusters are distributed within remyelinating white matter, we calculated a spline through the center of each lesion and determined the mean distance per cluster. We found that the core of the lesion at 7dpi, 12dpi and 18dpi was dominated by a microglia-derived signature: spatial cluster 13, which was enriched for antigen presentation genes *H2-Eb1*, *H2-Aa*, *H2-Ab1* and *Cd74*, cluster 6, enriched for *Top2a*, *Lgals3*, *Spp1* and *Gpnmb*, and cluster 2 which was enriched for *Csf1r* and *Lpl* (Extended Data Fig. 2c and 2e). Spatial cluster 10, matching OPC genes *Gpr17* and *Pdgfra*, was the next nearest cluster, followed by astrocyte cluster 0 (*Aqp4+*) which became more proximal to the repairing white matter core at the later phase of remyelination (18dpi) (Fig. 1i). Together, this reveals the complex architecture of repairing white matter, with multiple layers of glial populations in distinct reactive states.

### Monocyte-derived macrophages and lymphocytes sequentially enter the brain during remyelination

Infiltrating monocyte-derived cells are known to enter the brain after injury^34^ but the dynamics and transcriptional diversity of these cells during remyelination are unknown. We subclustered the putative monocyte-derived cells, identified by their expression of *Lyz2*, *Ms4a7* and *F13a1* (Fig. 1c) and lack of *Sall1*, a transcription factor specific to microglia^34–36^ (Extended Data Fig. 3a). Except for perivascular macrophages, all other monocyte-derived cells were essentially absent in saline controls (Extended Data Fig. 3b-c), demonstrating this population is exclusive to white matter damage and repair. This monocyte-derived population transiently entered the brain after demyelination at 3dpi but rapidly declined throughout remyelination (Fig. 2a). This population was unexpectedly diverse. We defined five distinct groups of these cells: the cluster MDM_1 expressed genes linked to phagocytosis^13^ (including *Spp1*, *Lgals1*, *Fabp5*, *Ctsd*, *Ctsb* and *Cd68*); MDM_2 specifically expressed the lysosomal proton pump *Atp6v0d2* in addition to other genes such as *Fbxo32*, *Adam33* and *Ccnd3*; the MDM_3 cluster was enriched for antigen presenting genes *H2-Eb1*, *H2-Ab1* and *Cd74,* in addition to the chemokine receptor *Ccr2*; a cluster of unclear function “Unknown” (including *Plxdc2*, *Nav3* and *Fat3*);and finally MDM_5, which expressed an interferon-responsive signature (including *Cxcl10, Ifit2* and *Ifit3*) (Fig. 2c-d, Extended Data Fig. 3d and Supplementary Table 5). The dendritic cell populations consisted of a conventional dendritic cell cluster cDC (expressing *Xcr1*, *Clec9a* and *Batf3*), plasmacytoid dendritic cell population pDC (expressing *Siglech*, *Ptprc* and *Tcf4*) and a small *Ccr7^+^* population of dendritic cells (migDC) that may be in the process of migrating from the brain parenchyma to lymph nodes^37^ (Fig. 2c-d). Finally, we identified perivascular macrophages (PVM), which specifically expressed *Cd163* and *Mrc1* marker genes^38^ (Fig. 2c-d and Extended Data Fig. 3b).

**Figure 2:**
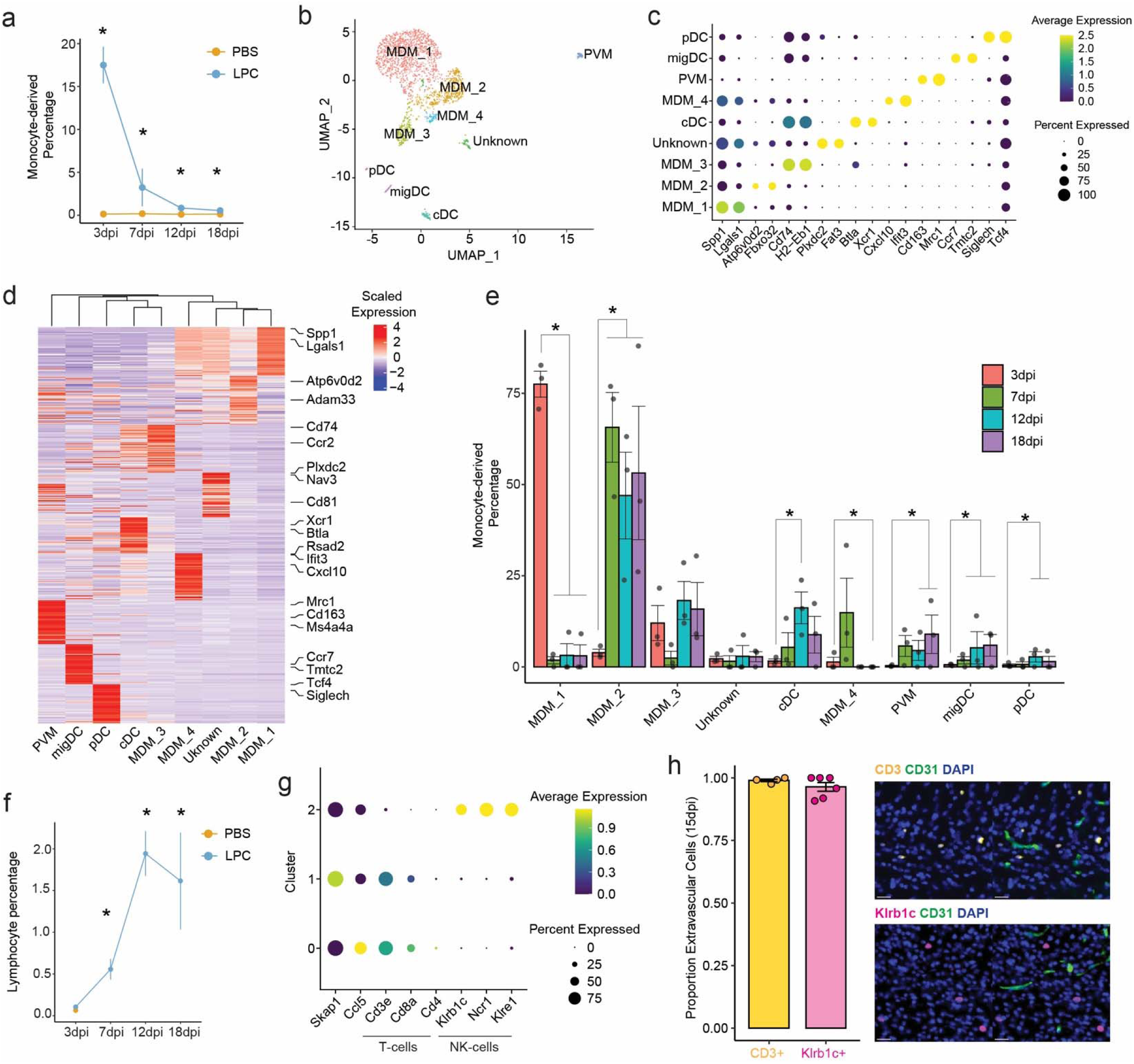
Sequential entry of monocyte-derived cells and lymphocytes during de-and remyelination. a) Percentage of monocyte-derived cells in the full snRNAseq dataset, plotted by PBS saline control and LPC. *=p-value<0.05. b) UMAP of monocyte-derived cells coloured by cluster. MDM=Monocyte-derived macrophages, cDC=conventional dendritic cells, pDC=plasmacytoid dendritic cells, migDC=migratory dendritic cells, PVM=perivascular macrophages. c) Dotplot of monocyte-derived clusters for selected marker genes. d) Heatmap of the top differentially expressed genes per monocyte-derived cluster. e) Percentage of monocyte-derived clusters per timepoint in LPC-injected animals only. Statistical significance was calculated by a differential abundance test (Methods). * = p-value<0.05. f) Percentage of lymphocytes in the full snRNAseq dataset, plotted by condition. * = p-value<0.05. g) Dotplot of cell type marker genes per lymphocyte cluster. h) (Left) Proportion of extravascular lymphocytes, T-cells (CD3^+^) and NK cells (KLRB1C^+^), defined by immunofluorescence. Extravascular cells defined as CD31 negative. (Right) Representative immunofluorescence images for T-cells (top) and NK cells (bottom).

We examined the temporal dynamics of these monocyte-derived subtypes during remyelination using a differential abundance test (Fig. 2e). The proportions of macrophages shifted dramatically, with MDM_1 representing the majority of cells at 3dpi (Fig. 2e), when monocyte-derived cells were most abundant (Fig. 2a), while MDM_2 dominated all subsequent timepoints (Fig. 2e). For rarer populations, we observed statistically significant gradual increases in proportions of dendritic cells including conventional (cDC), migrating (migDC), and plasmacytoid (pDC) (Fig. 2e). The proportion of Interferon-responsive macrophages (MDM_4) decreased over time while the percentage of perivascular macrophages increased (PVM) (Fig. 2e and Supplementary Table 6).

The diversity and temporal dynamics of lymphocytes are unknown during remyelination. In contrast to monocyte-derived cells, these cells were enriched in the remyelination phase, entering the repairing lesion at 7dpi and peaking at 12dpi and 18dpi, representing ∼2% of the total population (Fig. 2f). This population consisted of natural killer (NK) and T cells (Fig. 1c and 2g and Extended Data Fig. 3a-e). Unbiased analysis revealed three distinct clusters of lymphocytes, one of which was NK cells (cluster 2, positive for *Klrb1c, Ncr1* and *Klre1*), while both CD8^+^ and CD4^+^ T cells were distributed among clusters 0 and 1 (positive for *Cd3e* and either *Cd8a* or *Cd4*) (Fig. 2g and Extended Data Fig. 3b-e). To determine if these immune cells were truly in the brain, we performed co-staining with the vascular marker CD31. We observed that these cell types did not overlap with the vasculature, confirming these lymphocytes are truly parenchymal (Fig. 2h). Together, our dataset reveals how monocyte-derived macrophages and lymphocytes sequentially enter the damaged white matter during demyelination and repair respectively.

### Oligodendrocyte lineage cell dynamics reveals temporal dynamics and a novel OPC transcriptional state

We next focused on the oligodendrocyte lineage. We first examined the premyelinating oligodendrocyte population. This population consisted of OPCs (*Pdgfra*^+^) and pre-OLs (*Enpp6*^+^) but not *Aspa*^+^ mature myelinating oligodendrocytes^39^ (Fig. 3a-b and Extended Data Fig. 4a). Subclustering these cells revealed unexpected transcriptional diversity in control and remyelination conditions (Fig. 3c-d and Supplementary Table 7). We identified four clusters of OPCs (Fig. 3c): a large population of standard OPCs (OPC_1), two populations of cycling OPCs (OPC_4 and OPC_6, Extended Data Fig. 4b-c) and a previously undescribed population of *Glis3*^+^, *Col4a6*^+^ OPCs (OPC_5) (Fig. 3d and Extended Data Fig. 4d). We also identified two distinct subtypes of Pre-OLs, OPC_2 (positive for early myelination markers including *Casr*, *Mag* and *Mbp*) and OPC_3 (labelled by *Gpr17*^+^ and *Bmp4*^+^) (Fig. 3d). These OPC and pre-OL subtypes were dynamic throughout remyelination (Fig. 3e): In intact white matter, we observed a mix of standard and proliferating OPCs (OPC_1 and OPC_4) and both populations of pre-OLs. However, demyelination resulted in a dramatic decrease in standard OPC_1 at 3dpi, alongside a transient expansion of OPC_5, while the later phases of repair were characterized by increased proportion of pre-OLs relative to saline control (Fig. 3e and Supplementary Table 8).

**Figure 3:**
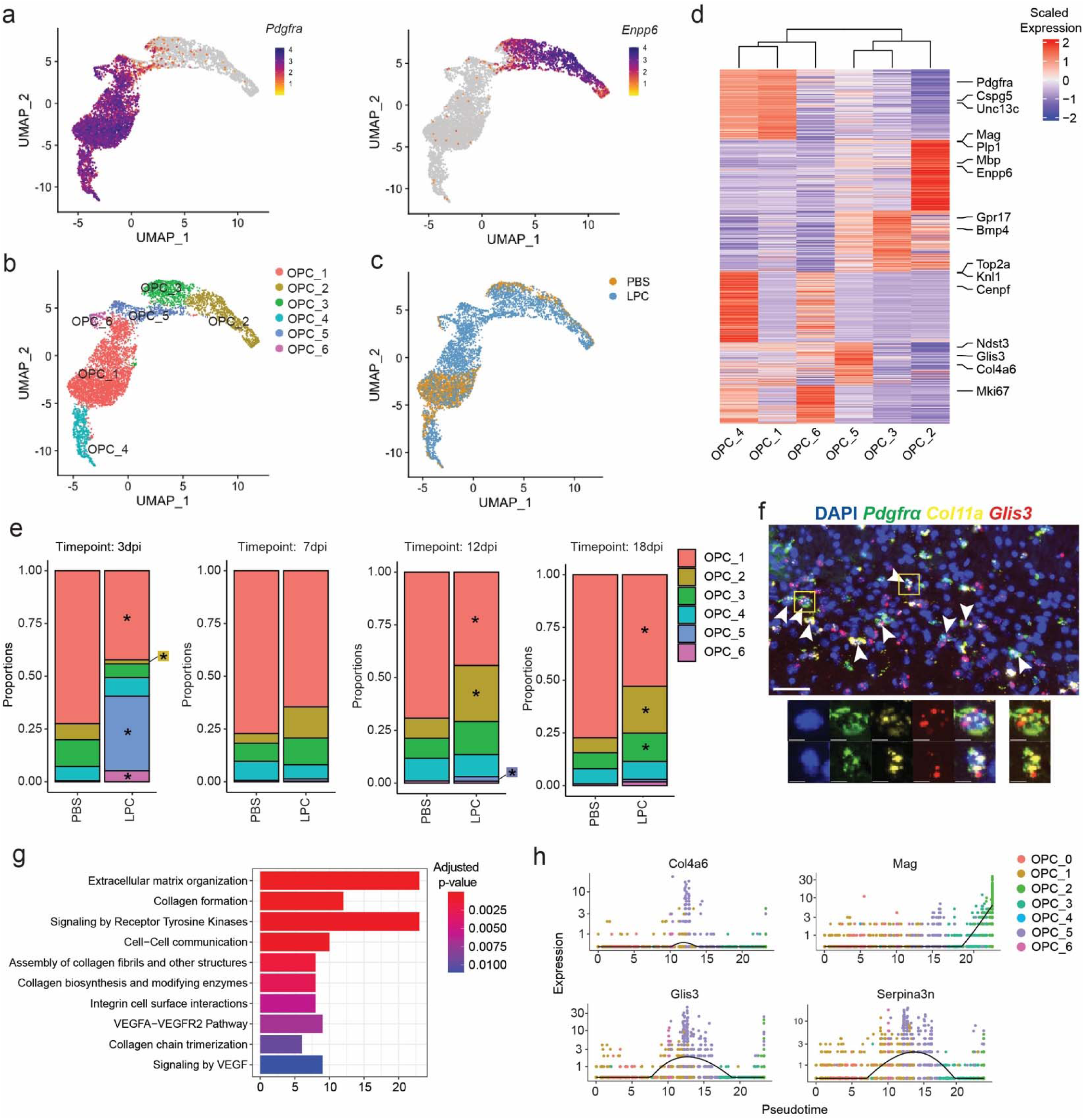
OPC dynamics reveal a demyelination-specific transitional state. a) UMAP of OPCs and pre-OLs with expression of *Pdgfra* (left) and *Enpp6* (right). b) UMAP of OPCs colored by cluster. c) UMAP of OPCs colored by condition. d) Heatmap of differentially expressed genes per cluster. e) Proportion of OPC and pre-OL clusters per timepoint and condition. *= p-value<0.05. Statistical significance determined by a differential abundance test. f) *In situ* hybridization of 3dpi LPC-injected lesions detecting *Glis3*, *Col11a* and *Pdgfra*. g) Gene ontology analysis of OPC_5 differentially expressed genes. h) Pseudotime analysis of OPCs during remyelination.

We further examined the *Glis3*+, *Col4a6*+ cluster (OPC_5), a transient, demyelination-specific OPC transcriptional state that has not been previously characterized. First, we confirmed the presence of this subtype in demyelinating lesions at 3dpi using *in situ* hybridization (Fig. 3f). In addition to the transcription factor *Glis3*, the OPC_5 signature was enriched for multiple components of the extracellular matrix, including collagens (*Col4a6*, *Col11a1, Col20a1, Col16a1, Col4a5* and *Col9a1*) and osteopontin (*Spp1*) (Extended Data Fig. 4e). Supporting this, a gene ontology analysis of enriched OPC_5 genes highlighted extracellular matrix organization, in addition to collagen biosynthesis and formation pathways (Fig. 3g). To test if OPC_5 represented a demyelination-specific intermediate state between OPCs and pre-OLs, we ordered OPCs from LPC-injected brains (all timepoints, 4,250 cells) on a pseudotime trajectory (Fig. 3h and Extended Data Fig. 4f) which indicated OPC_5 is a transitory state between OPCs and pre-OLs. We also demonstrated this state was not present in developmental myelination by performing a dataset integration and label transfer on a time course of corpus callosum myelinogenesis^39^ (Extended Data Fig. 4g-i). Together, this analysis revealed a transient, lesion-specific OPC state that precedes pre-OL formation and may modulate the extracellular matrix.

Within mature oligodendrocytes, we observed a dramatic shift in subtype composition during de-and remyelination (Extended Data Fig. 5a-b and Supplementary Tables 9-10). Two oligodendrocyte clusters (OL_0 and OL_1) predominated in saline-injected white matter, while four clusters were temporally dynamic throughout remyelination (Extended Data Fig. 5c-d). Three days after demyelination the majority of oligodendrocytes were OL_4, characterized by expression of *Klk6* and *Spp1* and a large increase in lipid metabolism genes (Extended Data Fig. 5e), which may represent an initial response to demyelination. In addition, exclusively at 3 dpi, we identified a small population of putatively injured or dying oligodendrocytes (OL_5) that specifically expressed heat shock proteins (*Hspa1a*, *Hspa1b* and *Hspb1*) and high levels of the ubiquitin-pathway genes *Ubb* and *Ubc* (Extended Data Fig. 5f). At later time points in remyelination, most mature oligodendrocytes were either characterized by an antigen presentation signature^16^ (OL_2, genes including *H2-D1*, *B2m* and *C4b*, see Extended Data Fig. 5g) or a *Pcdh7*^+^, *Man1a*^+^ subtype (OL_3, Extended Data Fig. 5h).

To cross-reference these transcriptional states with those identified in other damage contexts, we performed a gene set enrichment analysis on disease-associated oligodendrocyte populations identified in a large meta-analysis^18^ (Extended Data Fig. 5i), clear indicators of damaged white matter. This analysis demonstrated that OL_5 exhibits a disease-associated signature (“DA2”), while multiple oligodendrocyte states mapped to the “DA1” signature (Extended Data Fig. 5i). Interestingly, this analysis also identified an interferon-responsive “IFN” signature mapping to OL_2, and we subsequently identified a small subpopulation of interferon-responsive oligodendrocytes within OL_2, enriched for *Ifit3* and *Ifit2* (Extended Data Fig. 5j-i). This analysis revealed extensive transcriptional state diversity within all cells of the oligodendrocyte lineage throughout remyelination.

### Extensive microglial and astrocyte state diversity during remyelination

Microglia play distinct roles during de-and remyelination, phagocytosing debris and secreting factors to guide OPC differenciation^21–23^. We subclustered microglia (*Sall1*^+^, *Fcrls*^+^, *C1qa*^+^ cells, see Methods) and identified nine distinct microglial clusters, eight of which were exclusive to LPC injected animals (Fig. 4a-c). All microglial subtypes were defined by a large number of unique differentially expressed genes (Extended Data Fig. 6a and Supplementary Table 11) and we confirmed this striking heterogeneity using an alternative differential expression strategy^40^ (Extended Data Fig. 6b-c and Supplementary Table 12).

**Figure 4:**
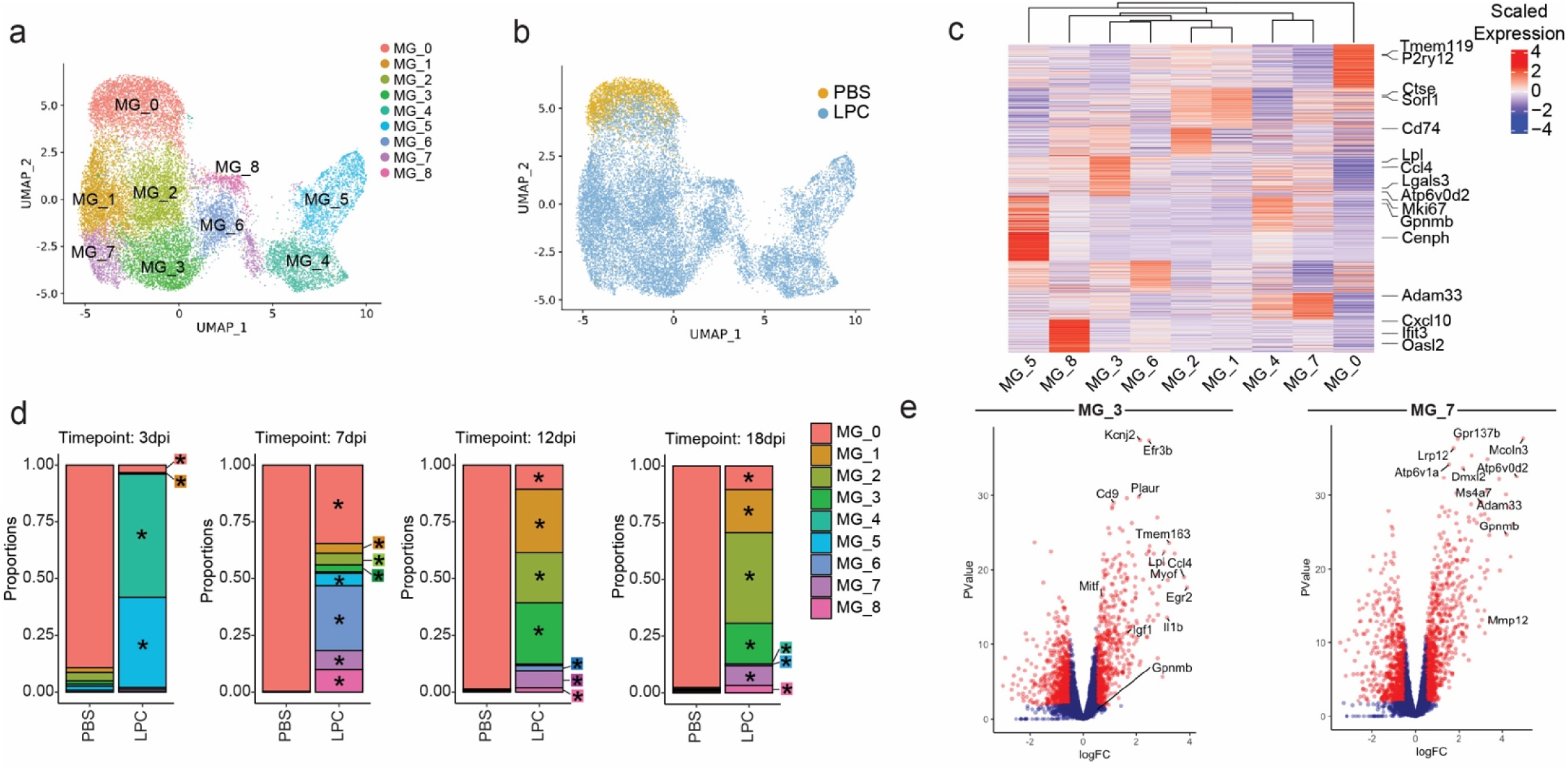
Extensive microglia heterogeneity throughout remyelination. a) UMAP of microglia plotted by cluster. b) UMAP of microglia plotted by condition. c) Heatmap of differentially expressed genes per microglia cluster. d) Proportion of microglial clusters per timepoint and condition. *= p-value<0.05. Statistical significance determined by a differential abundance test. e) Volcano plots two disease-associated microglial states, MG_3 and MG_7, demonstrating distinct enriched gene expression.

We identified several microglial states including a homeostatic signature (*Tmem119*^+^ and *P2ry12*^+^ MG_0), a *Ctse1*^+^ *Sorl1*^+^ cluster (MG_1), a proliferating cluster (*Mki67*^+^ and *Cenph*^+^ MG_5), a subtype characterized by an antigen presentation signature (*B2m*^+^, *H2-Ab1*^+^, *H2-D1*^+^ and *Cd74*^+^, MG_2) and a *Spp1*^+^, *Plin2*^+^ cluster enriched for purine nucleotide metabolism (MG_4, see Extended Data Fig. 6d) (Fig. 4a-c). In line with prior work in the cuprizone model of demyelination^20^, we also found two clusters resembling disease-associated microglia, MG_3 (*Lpl*^+^ and *Ccl4*^+^) and MG_7 (*Gpnmb*^+^, *Adam33*^+^ and *Atp6v0d2^+^*), each of which had notable differences in gene expression (Fig. 4e). Finally, we identified two clusters that exhibited differing levels of an Interferon-responsive gene signature, MG_6 and MG_8 (Fig. 4a-c). Both clusters were enriched for expression of *Stat1* and *Oasl2* while MG_8 additionally expressed *Cxcl10*, *Ifit2* and *Ifit3* (Fig. 4c), all of which are established markers of interferon signaling^41^. We validated these annotations by performing GSEA of microglial state signatures derived from a recent cross-species meta-analysis^42^, and the majority of these clusters mapped to known microglial states (Extended Data Fig. 6e).

We next examined the temporal dynamics of these microglial states from injury to repair (Fig. 4d and Supplementary Table 13). We observed a dramatic shift in microglia composition between conditions. At all timepoints, control microglia mostly consisted of the homeostatic white matter cluster MG_0. At 3 days after demyelination, microglia in LPC-injected brains were predominantly composed of proliferating MG_5 or *Spp1*^+^, *Plin2*^+^ MG_4. At 7dpi, we observed a striking degree of transcriptional diversification, identifying seven distinct microglial clusters including MG_4, MG_5, *Ctse1*^+^ *Sorl1*^+^ MG_1, antigen-presenting MG_2, disease-associated MG_3 and MG_7 in addition to interferon-responsive clusters MG_6 and MG_8. During the remyelination phase (12 and 18dpi), this diversity decreased, and microglia consisted mostly of clusters MG_1, MG_2 and MG_3.

Astrocytes also play crucial roles in remyelination^29,43^. We subclustered *Slca1a3*+ astrocytes^15^ and identified 8 distinct clusters (Fig. 5a-c, Extended Data Fig. 6f-g and Supplementary Table 14). Three of these clusters were significantly enriched in remyelination: AS_2, AS_5 and AS_6. At three days after demyelination, most astrocytes were either AS_6 or AS_2, strikingly different from saline injected controls (Fig. 5d and Supplementary Table 15). At later time points (12 and 18dpi), many astrocyte clusters were present, but AS_2 and AS_5 were statistically enriched in remyelination (Fig. 5d). These different clusters of astrocytes were defined by many unique genes: AS_6 expressed *Sorcs1*, *Ecrg4*, *Timp1* and proliferation markers *Mki67* and *Cdc20* (Fig. 5e). We validated the presence of AS_6 using *in situ* hybridization for *Timp1* at 3dpi (Fig. 5f). The AS_2 signature included *C3*, *Igfbp7* and *Cxcl10* while AS_5 profile included *Tmem108*, *Cxcl10* and *Ccl2* (Fig. 5g and Extended Data Fig. 6h). Notably, both AS_2 and AS_5 expressed genes characteristic of interferon-responsive signaling^15^ including *Cxcl10, Ifit2* and *Ifit3*. These results demonstrate extensive and persistent transcriptional heterogeneity of microglia and astrocytes during remyelination.

**Figure 5:**
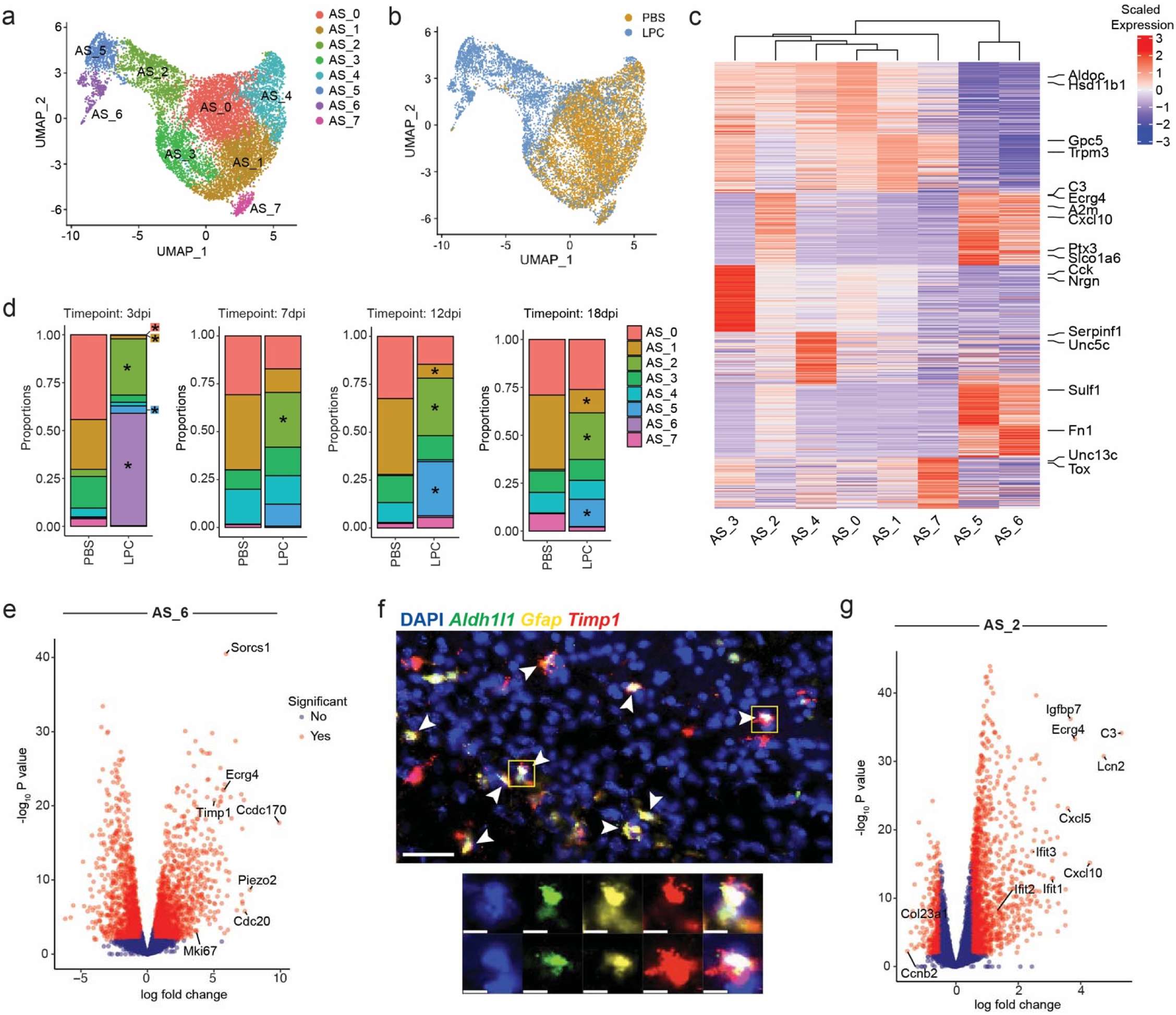
Astrocyte dynamics and heterogeneity during remyelination. a) UMAP of astrocytes, colored by cluster. b) UMAP of astrocytes colored by condition. c) Heatmap of the top differentially expressed genes per astrocyte cluster. d) Proportion of astrocyte clusters per timepoint and condition. *= p-value<0.05. Statistical significance determined by a differential abundance test. e) Volcano plot of remyelination-specific astrocyte cluster AS_6, showing differentially expressed genes relative to other clusters. f) *In situ* hybridization of 3dpi LPC-injected lesions detecting *Aldh1l1*, *Gfap* and *Timp1*. g) Volcano plot of remyelination-specific astrocyte cluster AS_2, showing differentially expressed genes relative to other clusters.

### Concerted type I interferon signaling drives the formation of a spatiotemporally regulated glial state

Across multiple cell types, including monocyte-derived macrophages, oligodendrocytes, microglia and astrocytes, we identified subclusters of cells expressing an interferon response gene signature during remyelination, suggesting this represents a distinct and coordinated transcriptional state across multiple glial cell types.

To investigate the nature of this multicellular program, we calculated an interferon signaling score to identify cells with strong expression of this signature (interferon-high). We found that interferon-high cells were only present in LPC injected animals, peaking at 7dpi (∼7.5% of total cells, Fig. 6a and Extended Data Fig. 7a), a shift that was mostly driven by microglia (Fig. 6b). This was further supported by an analysis of the raw module scores for the interferon-responsive gene signature (Extended Data Fig. 7b-c). These interferon-high cells mapped clearly to subpopulations identified in the unbiased clustering above, including in microglia (MG_6 and MG_8), astrocytes (AS_2 and AS_5), monocyte-derived macrophages (high in all clusters including MDM_4) and the small population of interferon-responsive oligodendrocytes (within OL_2) (Extended Data Fig. 7d-g).

**Figure 6:**
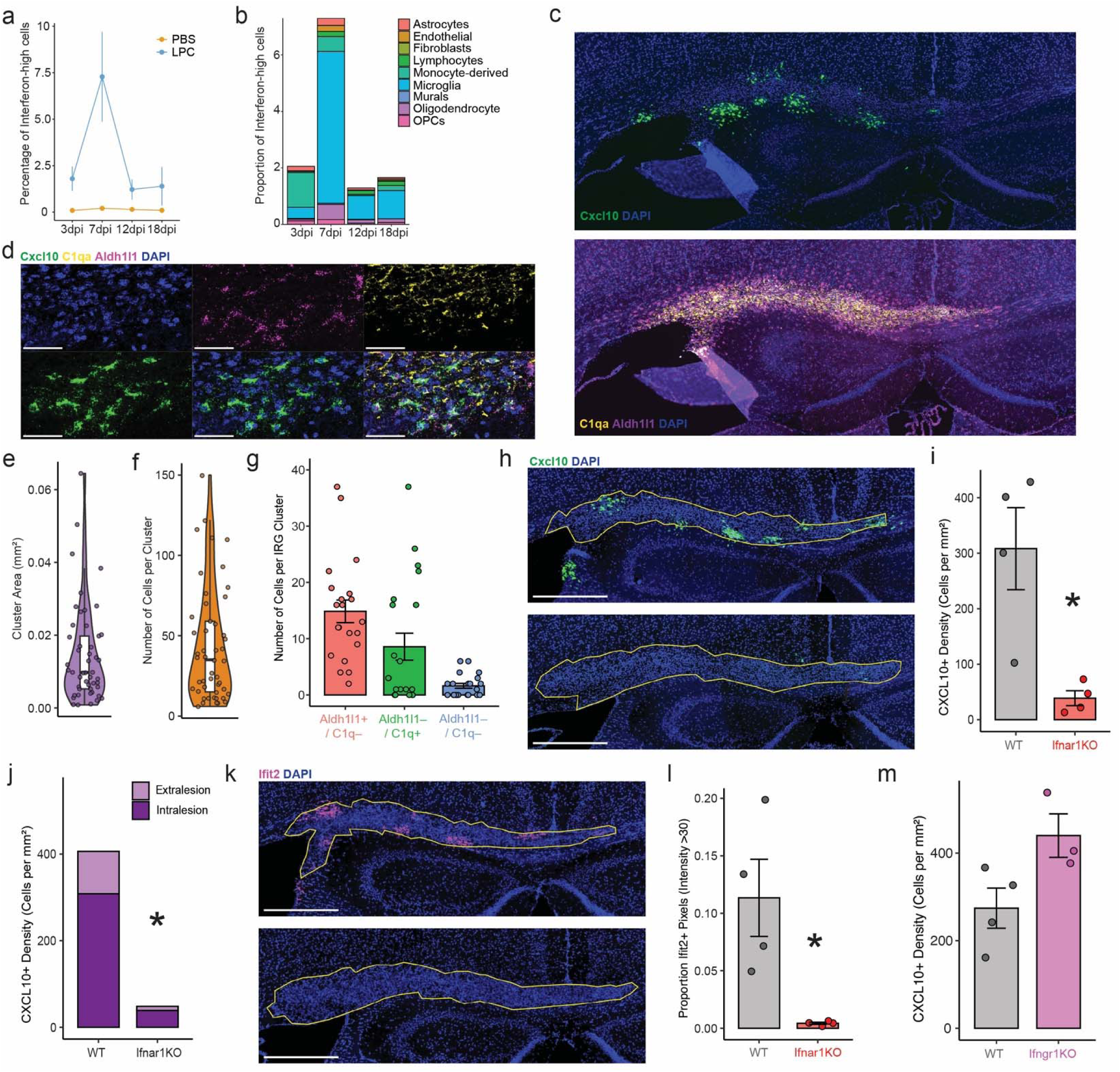
A concerted type I interferon response signature in glia prior to remyelination. a) Percentage of interferon-high cells, cells scoring above threshold for interferon responsive module score, in the full snRNAseq dataset. b) Interferon-high cells in the LPC condition, plotted by cell type per timepoint. c) Representative *in situ* hybridization for *Cxcl10* mRNA at 7dpi after LPC injection. (Top) Expression of Cxcl10. (Bottom) Expression of microglia and astrocyte cell type markers (*C1qa* and *Aldh1l1* respectively). d) High resolution confocal image of a *Cxcl10*^+^ IRG cluster with microglia and astrocyte cell type markers (*C1qa* and *Aldh1l1* respectively). e-g) IRG cluster area, cell number and cell type composition per 14um section (49 clusters from n=4 7dpi LPC-injected animals for area and density, 21 clusters from 4 7dpi LPC-injected animals for cell type composition). h) Representative image of *Cxcl10 in situ* hybridization at 7dpi in LPC-injected wild type (top) and *Ifnar1* knockout (bottom). i) Quantification of *Cxcl10*^+^ cell density per genotype. *= p-value<0.05. j) Breakdown of *Cxcl10*^+^ cell density as within (intralesion) and adjacent to (extralesion), the repairing white matter. *= p-value<0.05 k) Representative image of *Ifit2 in situ* hybridization at 7dpi in LPC-injected wild type (top) and *Ifnar1* knockout (bottom). i) Quantification of *Ifit2*^+^ cell density per genotype. *= p-value<0.05. m) Quantification of *Cxcl10* density across between wild-type and *Ifngr1* knockout animals at 7dpi.

Using differential expression, we examined the genes enriched in interferon-high cells and found robust expression of many interferon-response genes including *Cxcl10* and *Ifit2* (Supplementary Table 16). We termed these cells “interferon-responsive glia” (IRG), given that they were composed mostly of microglia, astrocytes and oligodendrocytes (Fig. 6b). To investigate the distribution of IRG during remyelination, we performed *in situ* hybridization targeting the IRG-specific gene *Cxcl10*. Strikingly, we observed that IRG formed clusters within and adjacent to demyelinated white matter (Fig. 6c and Extended Data Fig. 7h), with similar results for *Ifit2* (Extended Data Fig. 7i-j). These clusters were highly variable in both size and composition, with an average of 17582.4μm^2^ area and ∼50 cells per slice (Fig. 6e-f). A colocalization analysis using marker genes confirmed these clusters consisted mostly of astrocytes and microglia (Fig. 6g).

To identify the ligand that drives the formation of the IRG state in remyelination, we manipulated several receptors and pathways known to drive this signature^41^. We blocked type I and II interferon signaling using knockouts of *Ifnar1* and *Ifngr1* respectively. We found that deletion of *Ifnar1* attenuated the formation of *Cxcl10*^+^ IRG, both within and adjacent to the lesion (Fig. 6h-j). We confirmed this result with in situ hybridization for *Ifit2* (Fig. 6k), and with immunohistochemistry for the interferon signaling protein STAT1 (Fig. 6l). We observed no change in *Ifngr1* knockouts for any of these readouts (Fig. 6m and Extended Data Fig. 7k-i), nor did deletion of the cytosolic DNA-sensor *Sting1* impact STAT1 levels^44^ (Extended Data Fig. 7i), demonstrating that the formation of IRG is solely dependent on type I interferon signaling.

### IRG drive lymphocyte influx to modulate remyelination via CXCL10 signaling

To understand the function of the IRG state, we examined the specific marker gene *Cxcl10*, a cytokine that binds the receptor CXCR3 to mediate immune cell migration in a variety of organs^45^. We found that exposure of primary microglia and astrocytes to interferon-β drove robust secretion of CXCL10 protein into the media (Fig. 7a), confirming that these cell types can secrete CXCL10 in response to type I interferon stimulation. We also found that *Cxcr3* mRNA was specifically expressed in the infiltrating lymphocyte population (Fig. 7b).

**Figure 7:**
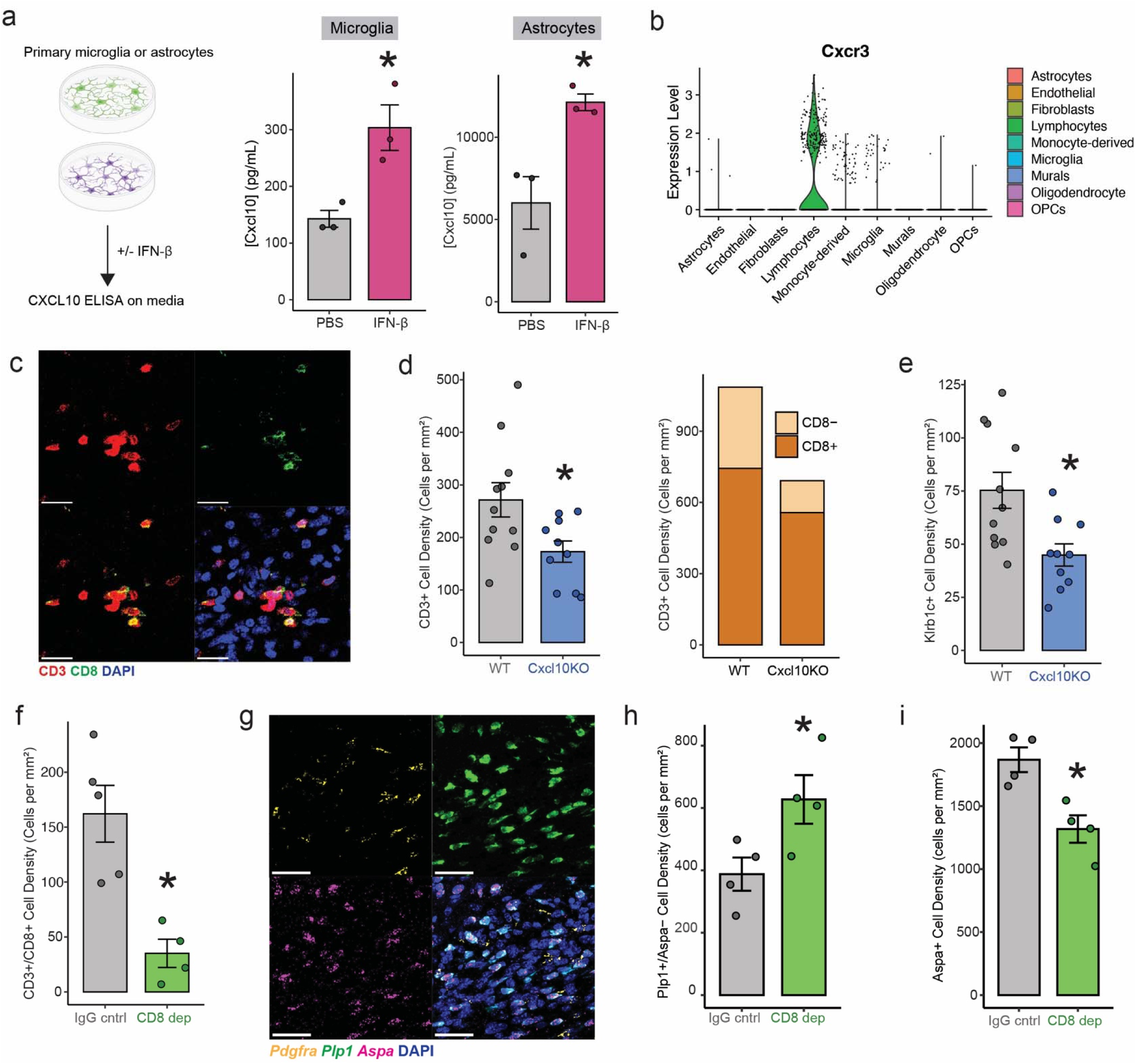
IRG drive lymphocyte brain infiltration via CXCL10 which in turn modulate oligodendrocyte differentiation. a) (Left) Schematic of primary glia ELISA for CXCL10 protein secretion. (Middle) ELISA for CXCL10 protein on media from primary microglia exposed to PBS or 25ng/ml murine Interferon-β. (Right) ELISA for CXCL10 protein on media from primary astrocytes exposed to PBS or 25ng/ml murine Interferon-β. b) Normalized expression of *Cxcr3* mRNA plotted per cell type in the full snRNAseq dataset. Each points represents a cell. c) Representative confocal immunofluorescence from a lesion showing detection of CD8 (CD3^+^, CD8^+^) and CD4 (CD3^+^, CD8^-^) T-cells. d) (Left) Density of T-cells in wild type and Cxcl10 knockout (LPC-injected animals at 15dpi). (Right) Breakdown of results by T-cell type. e) Density of NK cells in wild type and Cxcl10 knockout (LPC-injected animals at 15dpi). f) Density of CD8 T-cells in IgG control and CD8 antibody-injected animals (in LPC-injected animals at 18dpi). g) Representative image of *in situ* hybridization for oligodendrocyte lineage marker genes *Pdgfra*, *Plp1* and *Aspa*. h) Density of immature oligodendrocytes (*Plp1*^+^, *Aspa*^-^) across conditions. i) Density of mature oligodendrocytes (*Plp1*^+^, *Aspa*^+^) across conditions. For all panels *= p-value<0.05.

We hypothesized that IRG could drive T-and NK cell influx into the brain during white matter repair via CXCL10-CXCR3 signaling. To test this, we investigated immune cell infiltration during remyelination in *Cxcl10* knockout mice^46^. We first confirmed there was no major dysfunction of the peripheral immune system by examining the proportion of blood leukocytes using flow cytometry. We found no major differences in proportion across total CD45^+^ immune cells, NK-cells or total, CD8 and CD4 T-cells (Extended Data Fig. 8a). However, when we examined lymphocyte influx into repairing white matter, we found that *Cxcl10* knockout resulted in a decrease in T-and NK-cells entering the brain (Fig. 7c-e). Supporting this model, we found that attenuating the formation of IRG by deleting *Ifnar1* also resulted in the decreased brain infiltration of T-and NK cells during remyelination (Extended Data Fig. 8b). Together, this data demonstrates that, during remyelination, IRG coordinate neuroinflammation by driving lymphocyte influx.

To understand the functional impact of lymphocyte infiltration into remyelinating white matter, we developed a protocol for acute antibody-based depletion of CD8 T-cells (Extended Data Fig. 8c), the most abundant infiltrating lymphocyte cell type. We injected LPC and depleted CD8 T-cells throughout remyelination, acquiring tissue at 18dpi. To confirm successful CD8 T-cell depletion, we assayed the immune periphery by blood flow cytometry and found complete and specific ablation (Extended Data Fig. 8c-h). Critically, within the brain, we observed successful depletion of CD8 T-cells using immunostaining (Fig. 7f and Extended Data Fig. 8i-j).

To understand the impact of CD8 T-cells on remyelination, we focused on the oligodendrocyte lineage. We first used our snRNAseq data to develop an *in situ* hybridization panel to distinguish different stages of oligodendrocyte differentiation^39^ (Fig. 7g and Extended Data Fig. 8k). We found that CD8 T-cell depletion resulted in an increase in immature *Plp*1^+^, *Aspa*^-^; oligodendrocytes and a corresponding decrease in mature, myelinating *Plp1*^+^, *Aspa*^+^ oligodendrocytes (Fig. 7h-I and Extended Data Fig. 8l), while the number of *Pdgfra*^+^ OPCs were unchanged (Extended Data Fig. 8m), suggesting that CD8 T-cells guide oligodendrocyte maturation. Together, this demonstrates that IRG play a critical role in remyelination, driving lymphocyte infiltration via CXCL10-CXCR3 signaling, including CD8 T-cells, which in turn enhance oligodendrocyte differentiation.

### Human microglia and astrocytes adopt the IRG state and secrete CXCL10 in response to type I interferon

Finally, we examined if this interferon-response state could be induced in human glia. To model human IRG, we differentiated pluripotent stem cells into microglia^47,48^ (iMGs) or astrocytes^49^ (iAstros) and exposed these models to a low dose (25ng/ml) of recombinant human interferon-β. For iMGs, we performed single-cell RNA sequencing and found this stimulation drove robust formation of type I interferon-responsive microglia *in vitro* (Fig. 8a-c, Extended Data Fig. 10a-b) including upregulation of *CXCL10*, *IFIT2* and *IFIT3* (Fig. 8d, Extended Data Fig. 10c-e and Supplementary Table 17). We performed a similar experiment on iAstros, which were exposed to interferon-β and profiled using bulk RNA sequencing. Again, this revealed a strong upregulation of the type I interferon-response signature including *CXCL10*, *IFIT2* and *IFIT3* (Fig. 8e-f, Extended Data Fig. 9f and Supplementary Table 18). Critically, for both cell types we confirmed that interferon-β drove secretion of CXCL10 protein by performing ELISAs on conditioned media (Fig. 8g-h). Together, this demonstrates that human microglia and astrocytes can also adopt a CXCL10-secreting phenotype in response to type I interferon stimulation and provides a scalable model to examine additional functions of the human variant of this glial state.

**Figure 8:**
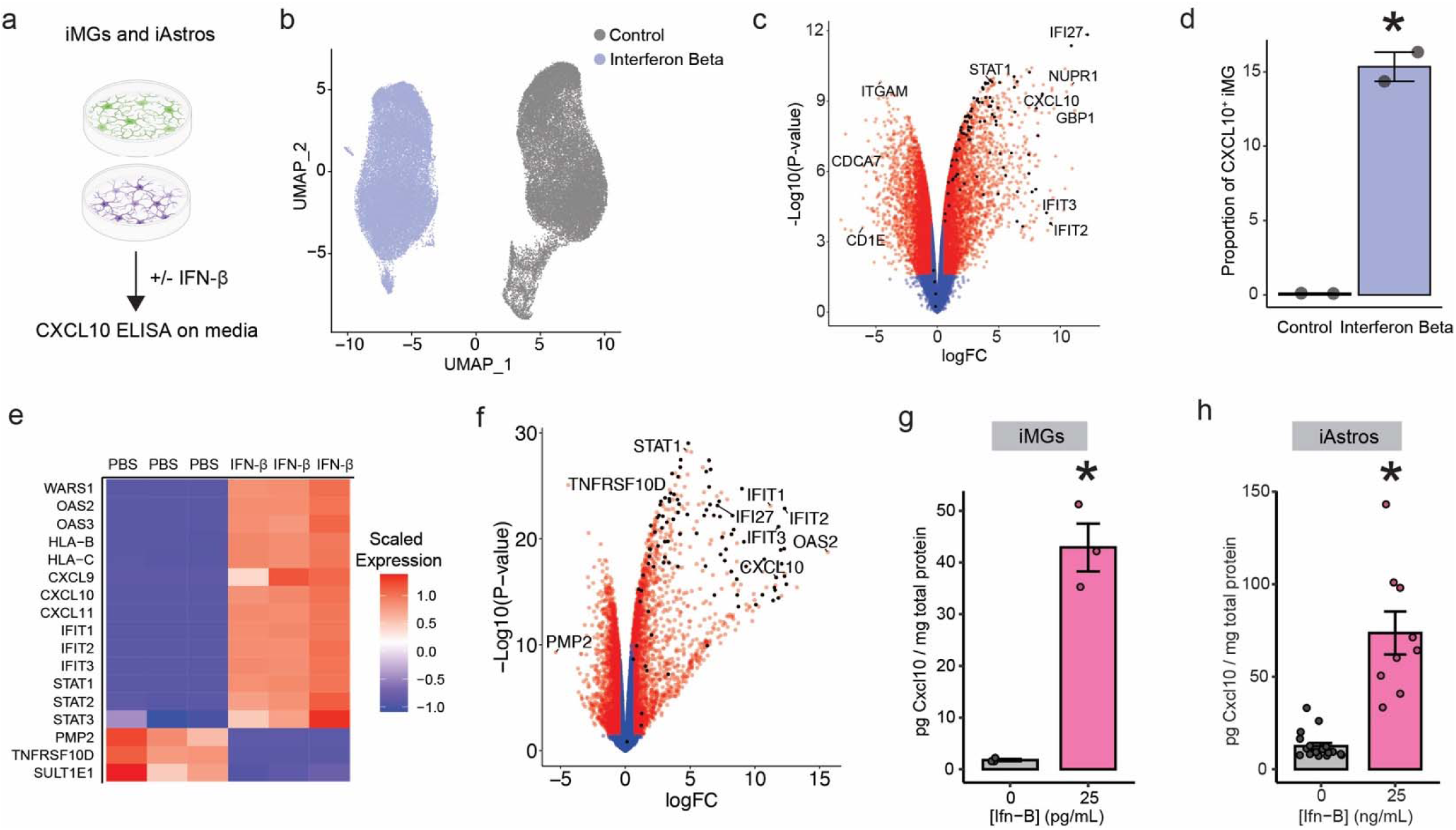
Induction of the IRG state in human stem cell-derived microglia and astrocytes. a) Schematic of experiment using stem cell-derived microglia (iMG) and astrocytes (iAstros). b) UMAP of iMG exposed to saline vehicle or 25ng/ml human interferon-β. c) Volcano plot highlighting differentially expressed genes in iMGs by exposure condition as determined by a pseudobulk analysis (Methods). Black dot indicates a type I interferon genes (from MSigDB). d) Proportion of iMGs positive for *CXCL10* mRNA (>1 mRNA) between conditions. e) Heatmap highlighting selected differentially expressed genes per iAstro sample (either PBS or IFN-β exposure). f) Volcano plot highlighting differentially expressed genes in iAstros by exposure condition. Black dot indicates a type I interferon genes (from MSigDB). g) ELISA for CXCL10 protein on media from iMG exposed to PBS or 25ng/ml human interferon-β. h) ELISA for CXCL10 protein on media from iAstros exposed to PBS or 25ng/ml human interferon-β. For all panels *= p-value<0.05.

## Discussion

Loss of myelin is a feature of aging, traumatic brain injury and numerous neurodegenerative diseases^5,7,50^. Remyelination is a critical reparative process that engages multiple cell types to repair denuded axons^1,2^. Here we provide an atlas of the transcriptional dynamics of repairing white matter, from initial injury to full repair. Our optimized nucleus extraction protocol enabled us to uncover extensive heterogeneity among all glial cell types, in addition to characterizing monocyte-derived macrophage dynamics and identifying a novel population of brain-infiltrating lymphocytes. Using spatial transcriptomics, we found that remyelinating white matter has a clear spatial structure, with a dense core of antigen presenting microglia and OPCs, surrounded by an outer rim of astrocytes.

A core feature of our dataset is the dense temporal sampling of cellular dynamics of both brain-resident glia and rare infiltrating immune cells. By tracking how these cell types and their transcriptional states shift over time, we made several important observations. Firstly, we found that peripheral cell types sequentially entered the brain at distinct timepoints: monocyte-derived macrophages were almost exclusive to the demyelination phase, while T-and NK cells infiltrated the lesion during remyelination. Given that blood-brain barrier disruption occurs early during demyelination^51^, this demonstrates the ability of the brain to summon distinct populations of immune cells in a temporally specific manner, likely using specific chemokines such as CXCL10. Of note, we did not identify regulatory T-cells which have been implicated in spinal cord remyelination^10^. This may reflect important regional differences^52^ or the need for additional timepoints to capture transient but critical brain-immune dynamics.

Our dense reconstruction of remyelination also enabled the discovery of a transient but coordinated interferon-response transcriptional state across microglia, astrocytes and oligodendrocytes, which we termed interferon-responsive glia (IRG). While this pathway is traditionally implicated in antiviral immunity^41^, interferon signaling has been identified during neurodevelopment, pathology and aging in both mouse and human brains^50,53,54^. However, the driving factors and downstream functions of this glial state are poorly understood. We demonstrated that the formation of IRG was dependent on type I interferon signaling, a family of ligands that includes the interferon-α gene family and interferon-β^41^. Functionally, we also found that IRG formed clusters in and around repairing white matter and secreted CXCL10 to drive the influx of T-and NK cells. This glial-immune interaction may represent a conserved response to brain injury. Indeed, we found that interferon-β could drive an interferon-responsive signature in human stem cell-derived microglia and astrocytes, including *CXCL10* mRNA expression and protein secretion. Further supporting this, a recent *in vitro* study of human glial-immune interactions which identified CXCL10 as a potent chemoattractant for T-cells in 3D neuronal cultures^55^.

To understand the downstream function of IRG clusters, we perturbed cytotoxic CD8 T-cells, the most abundant lymphocyte cell type that infiltrated the brain parenchyma during remyelination. We found that depletion of CD8 T-cells resulted in a decrease in the density of mature oligodendrocytes, suggesting that CD8 infiltration during repair enhances differentiation of the oligodendrocyte lineage. We hypothesize that this arises from an adaptive immune response, as we identified migratory dendritic cells^37^ in repairing white matter throughout de-and remyelination. While lymphocyte influx into the brain is traditionally associated with negative outcomes^50,56,57^, this is not always the case. Recent work has demonstrated that in models of axonal injury, CD4 T-cells play an important role in neuroprotection^58–60^. Decoding the activation status of these CD8 T-cells, their receptor clonality and their downstream cytotoxic target will be key to uncovering how peripheral immune infiltration can be beneficial in white matter repair. Moreover, advances in *ex vivo* T-cell manipulations mean this could be leveraged as a remyelination-enhancing cell therapy^60^. More generally, understanding how this adaptive immune response is beneficial in remyelination but deleterious in neurodegeneration will be an important step towards a context-specific understanding of brain-immune interactions.

## Acknowledgements

We thank all members of the Stevens and Macosko laboratories for helpful discussion when preparing the manuscript. We thank Tim Hammond for initial advice on the LPC model, Sahana Natarajan for providing primary glial cultures and Helena Barr for help with establishing peripheral immune cell flow cytometry. We thank the Broad’s Comparative Medicine, Genomic Platform and Flow Cytometry core facilities for their support and advice. This work was supported by the Alzheimer’s Association (ADSF-21-831114-C) and the International Neuroimmune Consortium, NIH Grant U19MH114821, The Stanley Family Foundation and the Howard Hughes Medical Institute. M.-J.D. was an Open Philanthropy Project Awardee of the Life Sciences Research Foundation. N.M. was supported by NIH training grant numbers: 5T32AG222-30 and 1F32AG079666-01. Graphical abstract and Figures 7a and 8b were created in Biorender.

## Contributions

Study design: M-JD, RR-S, EG, BS and EZM. Performed experiments: M-JD, RR-S, EG, NMN, JDixon with help from NM, CD, JDoman, JX and HJ. Performed analysis: M-JD, RR-S, SL, SG. Wrote the manuscript with input from all co-authors: M-JD, RR-S, BS and EZM. Provided reagents: CD. Contributed novel single-nucleus isolation methodology: NMN. Project supervision: BS and EZM.

## Supplementary Tables

Supplementary Table 1: Summary breakdown of cells per animal and cell type

Supplementary Table 2: Differential abundance by Cell type

Supplementary Table 3: Differential expression of spatial clusters from Slide-seqv2 data

Supplementary Table 4: Cell Type Module scores

Supplementary Table 5: Differential gene expression for monocyte-derived macrophages

Supplementary Table 6: Differential abundance for monocyte-derived macrophage

Supplementary Table 7: Differential gene expression for oligodendrocyte precursor cells

Supplementary Table 8: Differential abundance for oligodendrocyte precursor cells

Supplementary Table 9: Differential gene expression for oligodendrocytes

Supplementary Table 10: Differential abundance for oligodendrocytes

Supplementary Table 11: Differential gene expression for microglia (pseudobulk)

Supplementary Table 12: Differential gene expression for microglia (MAST)

Supplementary Table 13: Differential abundance for microglia

Supplementary Table 14: Differential gene expression for astrocytes

Supplementary Table 15: Differential abundance for astrocytes

Supplementary Table 16: Differential gene expression for “interferon-high” cells

Supplementary Table 17: Differential gene expression iMGLs exposed to interferon beta or control

Supplementary Table 18: Differential gene expression for iAstrocytes exposed to interferon beta or control

## Materials and Methods

### Methods - Experimental

#### Animal models

All experiments were performed C57BL/6J which was purchased directly from The Jackson Laboratory (cat# 000664). The following mice strains from The Jackson Laboratory were used: C57BL/6J (000664), B6(Cg)-Sting1tm1.2Camb/J (025805), B6(Cg)-Ifnar1tm1.2Ees/J (028288), B6.129S7-Ifngr1<tm1Agt>/J (003288), B6.129S4-Cxcl10tm1Adl/J (006087).

All mice were housed on a 12-h light/dark cycle between 68 °F and 79 °F and 30–70% humidity. All animal work was approved by the Broad’s Institutional Animal Care and Use Committee (IACUC).

#### Induction of remyelination using LPC

De-and remyelination was induced by injecting 1% lysophosphatidylcholine (LPC, #440154 EMD Millipore) into the mouse corpus callosum using stereotaxic surgery. As a vehicle, saline water was used. LPC was sonicated for 30min prior to surgeries. Animals were injected between P90-120 and this was kept consistent within experiments. Briefly, animals were anesthetized using isoflourane. Scalp was opened and a hole were drilled at:-1.2 anterior-posterior and 0.5 medio-lateral (distances calculated relative to bregma). The corpus callosum was injected at these coordinates and at a depth of-1.4 dorso-ventral to the skull. Injections were performed with a Nanoject III (Drummond) with a fire polished capillary. 500nl of LPC or control were injected at a rate of 1nl/sec for all mice used in the atlas, while 1000nl were injected at a rate of 2nl/sec for all mice used in functional studies. After injection, we waited 8 minutes before retracting the capillary to avoid back flow. Mice were sutured and returned to their home cage for recovery. During the surgery, mice received a dose of 5mg/kg meloxicam and 1mg/kg Buprenorphine SR.

#### Tissue Processing

Animals were processed at indicated days post injection (dpi) and brains were processed in two ways depending on downstream experiment. For snRNAseq or Slide-seqv2, mice were transcardially perfused with ice cold HBSS, brains rapidly extracted and frozen over the meniscus of liquid nitrogen using a small spoon. After freezing brains were stored in a sterile DNase/RNase free 5ml tube on a bed of frozen 1-2ml Optimal Cutting Temperature Medium (OCT, Sakura Biosciences). For the 3dpi brain for Slide-seqv2, the lesion made it difficult to mount on a Slide-seq bead array (see below). For this, the brain was frozen in an embedding cassette filled with OCT by holding the cassette above the meniscus of liquid nitrogen. Samples were kept on dry ice and immediately transferred to-80 degrees for storage.

For in situ hybridization (RNAscope) or Immunohistochemistry, mice were transcardially perfused with ice cold HBSS and brains were extracted and dropfixed in 25ml of 4% paraformaldehyde with gentle agitation at 4 °C overnight. The next day, brains were washed with ice cold PBS three times and sunk in 30% sucrose at 4 °C. After sinking, brains were immediately mounted in OCT within an embedding cassette using a combination of dry ice and 100% ethanol. Samples were kept on dry ice and immediately transferred to-80 °C for storage or - 20 °C for cyrosectioning. Samples were trimmed starting at the anterior face until the hippocampus was visible. After confirming presence of the lesion by staining a 14-micron coronal section with DAPI, further sections were taken and mounted on SuperFrost Plus^TM^ slides. Sections were stored at-80 ^O^C until further use.

#### Single-nucleus RNA sequencing of remyelinating lesions

##### Part I: Lesion identification and cryostat microdissection

Frozen mouse brains were microdissected in a cryostat (Leica). Nissel staining was performed as follows: After melting a tissue section onto a slide, the slide was air dried at room temperature and placed in 70% ethanol for 1 minute. Slides were washed in water and staining solution (Histogene™ Staining Solution, KIT0415, Thermofisher) was added for 4 minutes. Slides were washed in water and added to 70% ethanol, then 90% ethanol, then 100% ethanol for 30 seconds each. Finally slides were placed in 100% Xylene for 1 minute and mounted with a drop of permount. The region of demyelination was identified using Nissel staining to identify hippocampal landmarks and the characteristic hypercellularity associated with white matter damage and repair.

After finding the region of interest, the lesion was microdissected using a rectangular biopsy punch (generated by squeezing the tip of a 1mm Integra Miltex disposable biopsy punch, 33-31AA). Using a Keeler 4.5X headset for magnification, the biopsy punch pushed into lesioned white matter (based on location of nissel stain). The punch was then withdrawn and a 300um slice of frozen brain was cut by cryosectioning. Using cold, sterile forceps, lesions were microdissected from this slice and placed in a cooled DNase/RNase free PCR tube. Samples were stored at-80 for a maximum of 24 hours.

##### Part II: Nuclei extraction and FANS

Nucleus isolation was performed as described (https://www.protocols.io/view/frozen-tissue-nuclei-extraction-for-10xv3-snseq-rm7vz861xvx1/v2) with one modification: All washes and cell filtrations were performed in a 25ml falcon. After flow cytometry, nuclei were counted and loaded into the 10x Chromium V3.1 system. Reverse transcription and library generation were performed according to the manufacturer’s protocol.

#### Spatial transcriptomics with Slide-seqv2

Slide-seqv2 was performed as described (Stickels et al., 2021) (https://www.protocols.io/view/library-generation-using-slide-seqv2-81wgb7631vpk/v3). Prior to acquisition, the lesion was identified using the nissel protocol above (see *Part I: Lesion identification and cryostat microdissection).* Using a cryostat (Leica) 10um brain sections (acquired from fresh frozen brains) were melted onto Slide-seqv2 arrays. The lesion was aimed at the center of each array. After melting, bead arrays were immediately placed in a DNase/RNase free tube on dry ice for freezing. This enabled the collection of all samples to be processed in parallel. Processing occurred within 24 hours.

For processing, all reagents were diluted in ultrapure water (Life Technologies, Inc., 10977023), arrays were processed in parallel in Dnase/Rnase-free tubes. Arrays covered in tissue were transferred to 6xSSC (Life Technologies, Inc., 15557044) containing RNase inhibitor (Takara Bio, 2313B) and incubated for 15min. Arrays were then dipped in 1xRT buffer (Life Technologies, Inc., EP0753) and transferred to a premade Reverse Transcription reaction (Maxima RT: Life Technologies, Inc., EP0753; 10mM dNTP Life Technologies, Inc., 4303443; RNAase inhibitor: Takara Bio, 2313B; 50mM Template Switch Oligo: IDT, AAGCAG TGGTATCAACGCAGAGTGAATrG+GrG) for a 30min room temperature incubation. RT tubes were transferred to 52 C for a 90min incubation. Proteinase K and tissue clearing solution (Tris-HCl, pH 7.5: Life Technologies, Inc., 15567027; NaCl: American Bioanalytical, AB01915-01000; SDS (w/v): Life Technologies, Inc., 15553027; EDTA: Life Technologies, Inc., 15575020; Proteinase K: New England BioLabs, Inc., P8107S) was added to the RT tube and array was incubated at 37C for 30min. Beads were removed from glass and resuspended in TE-TW (TE buffer: Sigma-Aldrich, Inc., 8910-1L; Tween-20: VWR International, LLC, 100216-360) and subjected to x2 TE-TW washes, each followed by centrifugation for 2 min at 3000rcf to pellet the beads. Then, we exposed beads to 5min room temperature incubation in 0.1N NaOH (Sigma-Aldrich, Inc., followed by x2 TE-TW washes. After final spin, beads were incubated in second strand synthesis buffer (Maxima RT: Life Technologies, Inc., EP0753; 10mM dNTP Life Technologies, Inc., 4303443; dnSMRT oligo: IDT, AAGCAGTGGTAT CAACGCAGAGTGANNNGGNNNB; Klenow: New England BioLabs, Inc., M0212L) for 1 hour at 37C. Beads are then subjected to three TE-TW washes and loaded into WTA PCR (100 mM Truseq PCR primer: IDT, CTAC GACGCTCTTCCGATCT; 100 mM SMART PCR primer: IDT, AAGCAGTGGTATCAACGCAGAGT; Terra PCR mix: Takara Bio, 639284) with the following conditions: 98°C for 2min; 4 cycles of 98°C for 20sec, 65°C for 45sec, 72°C for 3min; 7 cycles of 98°C for 20sec, 67°C for 20sec, 72°C for 3min; and 72°C for 5min.

PCR clean-up was performed twice with 0.6x SPRI (Beckman Coulter, Inc., A63881) on cDNA libraries. Final and cDNA libraries were then examined on a bioanalyzer (Bioanalyzer High Sensitivity DNA kit: Agilent Technologies, Inc., 5067-4626) and Qubit (dsDNA high sensitivity kit: Life Technologies, Inc., Q32854) following manufacturer protocols. Tagmentation of 600pg of cDNA is performed according to Nextera DNA sample preparation manufacturer instructions (Illumina, Inc., FC-131-1096) using a Truseq-P5 hybrid constant oligo (IDT, AATG ATACGGCGACCACCGAGATCTACACTCTTTCCCTACACGACGCTCTTCCGATCT) and Nextera N7XX indexing primer (Illumina, Inc., FC-131-1001). Final libraries (4nM) were sequenced on a NovaSeq S4 with 150-200 million reads per sample at the Genomics Platform at the Broad Institute using the following read structure Read 1 42bp, Index 1 8bp, Read 2 41-60bp, and Index 2 0bp.

#### Immunohistochemistry

Brain slices mounted on SuperFrost Plus slides were brought to room temperature (RT) and rinsed in PBS to remove OCT. A hydrophobic barrier was drawn around the tissue using an ImmEdge® hydrophobic barrier pen (H-4000, Vector Laboratories). Slides were blocked in 4% normal goat serum (NGS) 0.3% Triton X-100 (A16046.AP, Thermofisher) in PBS at RT for 1 hour. Primary antibodies were applied overnight on a shaker at 4 ^O^C. After 3x 5-minute washes in PBS, secondary antibodies (at 1:800 dilution) were applied for 2 hours at RT. 4% NGS in PBS was used for both primary and secondary antibody solutions. Following application of secondaries, slides were again washed in PBS 3x times for 5 minutes each, then mounted with ProLong Gold Antifade Mountant with DNA Stain DAPI (P36935, Invitrogen) using VWR coverslips (48393-106), and sealed with clear nail polish (72180, Electron Microscopy Sciences). Stained slides were stored at 4°C until imaging. For epitopes that required citrate antigen retrieval (AR, see table below), after dissolving OCT, slides were incubated at 60 ^O^C for 90 minutes, then briefly rehydrated in PBS. Slides were then submerged in 1X pH 6.0 Citrate Buffer AR solution (C9999, Sigma-Aldrich) at 95°C for 20 minutes, then cooled in the buffer for an additional 20 minutes at RT. After AR, slides were briefly washed in PBS, then blocking proceeded as detailed above.

Table of all primary antibodies used:

| Antibody Target | Host Species | Cat Number, Provider | Dilution | Protocol |
| --- | --- | --- | --- | --- |
| STAT1 (D1K9Y) mAb | rabbit | 14994, Cell Signaling Technology | 1:250 | Citrate AR |
| CD3 | hamster | 553238 BD Bioscience | 1:400 |  |
| CD8a | rat | YTS105.18, Biorad | 1:200 |  |
| CD4 | rat | 100506, Biolegend | 1:400 |  |
| CD31 | rat | DIA-310, Dianova | 1:20 | Both with and without Citrate AR |
| NK1.1/CD161c/ KLRB1c | rabbit | 39197, Cell Signaling Technology | 1:400 | Citrate AR |

The following secondary antibodies were used:

Goat anti-rabbit IgG (H+L) Alexa Fluor 568 (A-11011, Thermofisher), Goat anti-rabbit IgG (H+L) Alexa Fluor 647 (A-21245, Thermofisher), Goat anti-rat IgG (H+L) Alexa Fluor™ 568 (A-11077, Thermofisher), Goat anti-rat IgG (H+L) Alexa Fluor 647 (A-21247, Thermofisher), Goat anti-Mouse IgG2b Alexa Fluor 647 (A-21242, Thermofisher), Goat anti-Syrian Hamster IgG (H+L) Alexa Fluor 647 (A-21451, Thermofisher), and Goat anti-Chicken IgY (H+L) Alexa Fluor 647 (A-21449, Thermofisher).

#### *In situ* hybridization

Mouse brain slices mounted on SuperFrost Plus slides were brought to room temperature (RT) and rinsed in PBS for 5 minutes to remove OCT. Subsequently, slides were incubated at 60^°^C for 30 minutes, then postfixed in 4% PFA for 15 minutes on ice. Slides were then dehydrated, and pretreated according to the manufacturer’s (Advanced Cell Diagnostics) instructions for sample preparation of fixed frozen tissue for the RNAScope Multiplex Fluorescent Assay v2, with the following modifications: Brain sections were treated with RNAScope Hydrogen Peroxide for 10 minutes at RT following tissue dehydration. Slides were incubated in RNAScope Target Retrieval solution for 10 minutes at 90°C.

After sample preparation, the RNAScope Multiplex Fluorescent v2 Assay (323100) was performed according to the manufacturer’s instructions. Stained slides were stored at 4°C until imaging.

The following probes from Advanced Cell Diagnostics were used: Mm-Glis3 (314161), Mm-Col11a1-C2 (439241-C2), Mm-Pdgfra-C3 (480661-C3), Mm-Timp1 (316841), Mm-Gfap-C2 (313211-C2), Mm-Aldh1l1-C3 (405891-C3), Mm-Aspa (425891), Mm-C1qa-C2 (441221-C2), Mm-Cxcl10-C3 (408921-C3), Mm-Ifit2 (577211), Mm-Flt1 (415541), and Mm-Plp1-C2 (428181-C2). The following dyes from Akoya Biosciences were used: TSA Plus Fluorescein (SKU NEL741001KT), TSA Plus Cyanine 3 (SKU NEL744001KT), and TSA Plus Cyanine 5 (SKU NEL745001KT).

### Imaging

All imaging was performed with either a Zeiss LSM 900 or a Zeiss Axioscan 7 Slide Confocal imaging was acquired using a 20x or 63x oil objective. Whole sections were imaged with a Zeiss Axioscan 7 Slidescanner using exposure and intensity settings that were constant across samples within technical replicates. Scanner using a 20x objective.

### Flow cytometry on peripheral immune cells

Blood isolations were carried out prior to performing intracardiac perfusions (see above). A sterile insulin syringe with fixed needle (Air-Tite, cat. #EU05308) was pre-filled with 50ul FACS buffer (HBSS with 0.5% BSA and 2mM EDTA) and stored on ice. Similarly, 10ml of FACS buffer was added to 15ml conical tubes and placed on ice. Blood was collected by puncturing the right atrium with syringe, flushed into 15ml conical tubes, vortexed for 5 seconds and returned to ice. Blood cells were pelleted by centrifugation (500g for 5 minutes at 4 degrees) and red blood cells were lysed with ACK Lysis Buffer (A10492-01, ThermoFisher). Immune cells were resuspended in FACS buffer and washed once. Cells were spun down (500g for 5 minutes at 4 degrees), resuspended in 50ul Fc block and 1/1000 viability stain (Fixable Viability Dye eFluor™ 780, Biolegend) and incubated on ice for 10 minutes. A staining solution, consisting of PE/Cyanine7 anti-mouse CD45 Antibody (1/200, 103113, Biolegend), Alexa Fluor® 488 anti-mouse NK-1.1 Antibody (1/50, 108717, Biolegend), Brilliant Violet 421™ anti-mouse CD4 Antibody (1/50, 100437, Biolegend), PE anti-mouse CD8a Antibody (1/50, 100707, Biolegend) and APC anti-mouse CD3 Antibody (1/50, 100235, Biolegend) was added and cells were incubated for another 20 minutes on ice. FACS buffer was added and cells were washed twice. After a third spin, cells were resuspended in 100ul ICC Fixation Buffer (cat. #550010, BD) and incubated for 20 minutes at room temperature. 100ul FACS buffer was added and cells were washed two more times prior to flow cytometry. Flow cytometry was performed on a CytoFLEX LX and analyzed with CytExpert v2.4.0.28.

### Primary microglia and astrocyte culture and treatment

Freshly dissected brains from P1-P5 pups were pooled in a 15mL tube with HBSS (2-3 brains per tube). HBSS was then replaced with 10mL of plating media (DMEM (10569-010, Gibco) with 1% penicillin-streptomycin and 10% heat inactivated FBS (10082147, Life Technologies)). Brains were subsequently triturated with a 10mL serological pipette, 1000μL pipette, and then a fire-polished glass Pasteur pipette. The solution was allowed to settle for 1 minute, then the supernatant was transferred to a 50mL tube and centrifuged for 5 minutes at 300g. The supernatant was discarded, and the pellet resuspended in plating media (2.5mL per brain). 2.5mL of resuspended cells were added to a T75 flask containing 15mL prewarmed plating media. After a 24-hour incubation at 37°C, media was removed, cells were washed with 10mL prewarmed HBSS, and 15mL prewarmed plating media was added.

On day 6-8 *in vitro*, microglia were removed by shaking flasks for 1 hour at 200 RPM in an incubator at 37°C, then the media was replaced with 20mL prewarmed plating media. Flasks were subsequently incubated at 37°C overnight on a shaker at 240 RPM to remove OPCs, transferred to the upright position, and washed twice with 10mL prewarmed HBSS while shaking vigorously to dislodge any remaining OPCs. Media was replaced with 15 mL prewarmed plating media, then microglia were harvested on *in vitro* day 10-13.

To collect remaining microglia, flasks were incubated for 30-60 minutes at 37°C on a shaker at 200 RPM, then media was collected in 50mL tubes and centrifuged for 5 minutes at 300g. After carefully removing the supernatant, the pellet was resuspended in 1mL NbActiv4 media (NB4, Transnetyx) with 40ng/mL mouse macrophage colony stimulating factor (mCSF1) (Peprotech). Cell concentration was determined with a Countess 3 Automated Cell Counter (AMQAX2000, Invitrogen), then cells were plated in a 24 well plate at 10^5^ cells / well. 1mL NbActiv4 media with 40ng/mL M-CSF was added to each well. Media was replaced every 3 days after plating.

After harvesting the microglia, 20mL of prewarmed plating medium was added to the flasks, then flasks were incubated overnight at 37°C on a shaker at 240 RPM. Flasks were subsequently transferred to the upright position, washed twice with 10mL prewarmed PBS while shaking vigorously, and incubated with 5mL TrypLE Express (12605036, Thermofisher) for 20 minutes at 37°C to dislodge the adherent astrocytes. 20mL NbActiv4 with 1% penicillin-streptomycin was added, and the cell suspension was centrifuged for 5 minutes at 300g. After removing the supernatant, cells were resuspended in 1mL plating medium, counted with a Countess™ 3 Automated Cell Counter, and plated in a 24 well plate at 5×10^4^ cells/well with 0.5mL plating medium per well. UV sterilized poly-D-lysine coated coverslips were placed in each well prior to plating. Media was replaced every 3 days after plating.

For cell treatment: mouse IFN-β (8234-MB-010/CF, R&D Systems) or equivalent volume of PBS vehicle were added to each well for a final concentration of 25ng/mL Interferon. Media and cells were collected after 24 hours incubation at 37 ^O^C. Media aliquots were flash frozen on dry ice and stored at-80°C until use in ELISAs (see below).

### Induced Microglia (iMG) differentiation

H1 embryonic stem cells (WiCell) were used. Differentiation of microglia from induced pluripotent stem cells (iPSCs) was performed as described(Abud et al. 2017). Briefly, ESC were cultured in Essential 8 (E8) (Thermo Fisher Scientific) media on Matrigel (Corning) coated 6-well plates. When confluent, cells were dissociated with Accutase (Stem Cell technologies). 200,000 cells/well were resuspended in E8 containing 10uM Y27632 ROCK inhibitor (Selleckchem) in a low adherence 6-well plate (Corning).

For the first 10 days, cells were cultured in HPC medium [50% IMDM Thermo Fisher Scientific), 50% F12 (Thermo Fisher Scientific), ITSG-X 2% v/v (Thermo Fisher Scientific), L-ascorbic acid 2-Phosphate (64 ug/ml, Sigma), monothioglycerol (400mM, Sigma), Poly(vinyl) alcohol (PVA) (10mg/ml, Sigma), Glutamax (1X, Thermo Fisher Scientific), chemically-defined lipid concentrate (1X, Thermo Fisher Scientific) and non-essential amino acids (Thermo Fisher Scientific)]. At day 0, embryoid bodies (EB) were gently collected, centrifuged at 100x*g* and resuspended in HPC medium supplemented with 1uM ROCK inhibitor, FGF2 (50 ng/ml, Thermo Fisher Scientific), BMP4 (50ng/ml, Thermo Fisher Scientific), Activin-A (12.5ng/ml, Thermo Fisher Scientific) and LiCL (2 mM, Sigma), then incubated at in hypoxic incubator (5% O2, 5% CO2, 37°C). On day 2, cells were gently collected and the media changed to HPC medium supplemented with FGF2 (50ng/ml, Thermo Fisher Scientific) and VEGF (50 ng/ml, PeproTech) and returned to the hypoxic incubator. On day 4 cells were gently collected and media changed to HPC medium supplemented with FGF2 (50ng/ml, Thermo Fisher Scientific), VEGF (50 ng/ml, PeproTech), TPO (50 ng/ml, PeproTech), SCF (10ng/ml, Thermo Fisher Scientific), IL6 (50ng/ml, PeproTech) and IL3 (10ng/ml, PeproTech) and incubated in normoxic incubator (20% O2, 5% CO2, 37°C). At day 6 and 8, 1 ml of day 4 media was added in each well. On day 10, cells were collected, counted using trypan blue and frozen in Cryostor (Sigma Aldrich) in aliquots of 300,000-500,000 cells.

To start iMG differentiation, cells were thawed, washed 1x with PBS and plated at 100,000-200,000 cells per well in 6-well plate coated with matrigel in iMG media [(DMEM/F12 (Thermo Fisher Scientific), ITS-G (2% v/v, Thermo Fisher Scientific), B27 (2% v/v, Thermo Fisher Scientific), N2 (0.5% v/v, Thermo Fisher Scientific), monothioglycerol (200 mM, Sigma), Glutamax (1X, Thermo Fisher Scientific), non-essential amino acids (1X, Thermo Fisher Scientific)] supplemented with M-CSF (25 ng/ml, PeproTech), IL-34 (10 ng/ml, PeproTech) and TGFB-1 (50ng/ml, PeproTech). Cells were fed every 2 days and replated at day 22. On day 30, cells were collected and replated in iMGL media supplemented with M-CSF (25 ng/ml, PeproTech), IL-34 (10 ng/ml, PeproTech), TGFB-1 (50ng/ml, PeproTech), CD200 (100ng/ml, VWR) and CX3CL1 (100ng/ml, PeproTech) to a final concentration of 40,000 cells/cm^2^. Cells were used at day 40-45.

### Induced Astrocyte differentiation (iAstros)

To generate iAstrocytes, H1 human embryonic stem cells (hESCs) were first electroporated with “AAVS1 HA L-SA-NeoR*-TRE3G-NFIB-P2A-SOX9-IRES-ZeoR-AAVS1 HA-R” (“pGEP-AAVS1-BS9”), AAVS1 TALEN L, and AAVS1 TALEN R vectors to knock-in human NFIB and SOX9 transcription factors into the AAVS1 genomic safe harbor locus. hESCs were dissociated with accutase and 2e6 cells were electroporated with 10 μg of donor plasmid and 1.5 μg of each TALEN vector using a Neon Transfection System (1050 V, 30 ms, 2 pulses). hESCs were cultured in E8 media with CloneR2 for 3 days and then switched to E8 media with 100 μg/mL geneticin to select for genomic integration of doxycycline-inducible NFIB-SOX9-IRES-ZeoR. After expansion, H1 cells were split with accutase and seeded at 5,200 cells / cm^2^ on growth factor-reduced matrigel plates in E8 with rock inhibitor. iAstrocytes were generated using a published differentiation media protocol(Canals et al. 2018). Briefly on DIV0, hESCs were cultured in E8 media with 500 ng/mL doxycycline to initiate NFIB-SOX9 expression. On DIV1 and DIV2 iAstrocytes were cultured in expansion media (EM: DMEM/F12, 10% heat-inactivated FBS, 1% N2, 1% Glutamax, 500 ng/mL doxycycline) with 200 μg/mL zeocin. From DIV3-5, iAstrocytes were cultured in 3:1, 1:1, and 1:3 ratios of EM and FGF media (FGF: Neurobasal, 2% B27, 1% NEAA, 1% Glutamax, 1% HI-FBS, 500 ng/mL doxycycline, with fresh 10 ng/mL BMP4, 5 ng/mL CNTF, 8 ng/mL FGF). iAstrocytes were cultured in FGF media on DIV6 and dissociated on DIV7 with accutase before freezing in CryoStor CS10. DIV7 iAstrocytes were thawed onto matrigel-coated plates in FGF media and given a full media change on DIV8. On DIV10, iAstrocytes were switched to maturation media (MM: 50:50 Neurobasal:DMEM/F12, 1% N2, 1% Sodium Pyruvate, 1% Glutamax, 5 ng/mL heparin-binding EGF, 5 μg/mL N-acetyl-L-cysteine, 500 ng/mL doxycycline) with fresh 10 ng/mL BMP4, 10 ng/mL CNTF, 500 μg/mL dbcAMP. iAstrocytes were cultured in MM with ½ media changes every other day until DIV20. iAstrocyte conditioned media was centrifuged at 1000g for 10 minutes and stored at-80°C.

### Interferon Exposure to iMG and iAstros

Human IFN-β (300-02BC, Peprotech) or equivalent volume of PBS vehicle were added to each well for a final concentration of 25ng/mL. Media and cells were collected after 24 hours incubation at 37°C. Media aliquots were flash frozen on dry ice and stored at-80 ^O^C until use in ELISAs (see below).

### ELISAs

ELISAs for mouse (DY466-05, R&D Systems) and human (DY266, R&D Systems) CXCL10 were performed according to the manufacturer’s instructions. Prior to the assay, media from primary mouse astrocytes, primary mouse microglia, human iPSC-derived astrocytes (iAstros), or human iPSC derived microglia (iMG) were diluted 1:1 to 1:100 with sterile PBS to ensure sample CXCL10 concentrations fell within the standard curve. Total protein concentrations were determined using a Pierce™ BCA Protein Assay Kit (23225, Thermofisher). Absorbances were read with a Perkin Elmer EnVision 2104 Multilabel Plate Reader.

### Bulk RNA sequencing

For iAstro bulk RNA sequencing, media was removed and adherent iAstros were incubated with 5mL of TrypLE Express for 10 minutes at 37 ^O^C. Cells were collected and cell suspension was centrifuged in a 15mL tube for 5 minutes at 300g. The supernatant was discarded, then the cells were resuspended in 10mL PBS and centrifuged again for 5 minutes at 300g. After discarding the supernatant, the cells transferred to a fresh microcentrifuge tube. These were resuspended in 1mL PBS and washed again. Once PBS is removed, the pellet, was flash frozen on dry ice, and stored at-80°C until RNA extraction for bulk transcriptomics. RNA was extracted using the RNeasy Mini kit (Qiagen) and bulk RNA-sequencing was performed by Novogene Inc, using directional mRNA library preparation (Poly A enrichment) and each sample was sequenced at a depth of 30M reads.

## Methods - Analysis

### Alignment, preprocessing and clustering of single-nucleus data

For single-nucleus RNAseq of white matter: Cellranger (version 6, 10X Genomics) was used for demultiplexing, barcode preprocessing, generation of fastq files, alignment and the counting of unique molecular identifiers (UMI). Please note that single-cell RNAseq of iMGs is dealt with separately below.

All downstream analysis was performed using R version 4 or 3.6.3, using the Seurat package (v4) (Stuart et al. 2019). All data and metadata (replicate and condition) were incorporated into a single object and the percentage of mitochondrial RNA per cell was calculated. We only considered nuclei with the following characteristics: 1) Number of Genes: 750-6000, 2) Number of UMIs: 1500-25000, 3) Percentage mitochondrial RNA <1%. This yielded 131,521 single nuclei consisting of glial and neuronal cell types. Neurons were considered to be microdissection artefacts and removed prior to final clustering of 77829 glial-immune nuclei. Neurons were identified by expression of *Snap25*, *Rbfox3*, *Gad1* and *Gad2*. After filtering the neurons, cell types were identified using marker genes (Fig. 1c) and subclustered for analyses. Doublets were identified as expressing canonical marker genes of two or more cell types and removed.

Each cell type was subclustered: cells of type were normalized using the NormalizeData function in Seurat with the LogNormalize option and a 10,000 scale factor. Cells were scaled and clusters were identified based on gene expression using Louvain method. This step was performed twice, the first time to identify doublet clusters sharing 1 or more key cell type markers, the second time these doublets were filtered out prior to normalization. For each cell type, cluster validation was performed by identifying unique differentially expressed genes (>50 unique DEGs required per cluster). Additional details for specific cell types:

#### Oligodendrocyte Precursor Cell (OPC) clustering

OPCs were subclustered based on expression of well-established marker genes (*Pdgfra* and *Vcan*). These cells underwent an initial round of clustering to identify 234 *Plp1*+ oligodendrocyte/OPC doublets which were removed and analysis repeated (top 20 principal components used for SNN analysis and clustering resolution of 0.3).

This analysis identified 7 different OPC subtypes. We performed a differential analysis using MAST and determined using upset plots that two of these states were highly similar. We merged these two clusters into final cluster 1. All other clusters remain the same from this analysis.

### Differential Abundance

All proportional data plotted in figures is untransformed. However, for the calculation of p-values, this data underwent a logit transformation prior to running differential abundance tests (using propellor from the R library, “speckle” version 0.99) (Phipson et al. 2022).

### Single-nucleus RNA seq Differential Expression analysis

For most cell type specific differential expression analysis, we used a pseudobulk approach using “edgeR” (v4.2) (Robinson, McCarthy, and Smyth 2010). Briefly, raw counts were summed by cluster and sample (ie. animal) using the Seurat2PB function in edgeR. Sample clusters with a library size < 5e4 were filtered out as were lowly expressed genes (using filterByExpr function in edgeR). We performed a TMM normalization, dispersion estimation and differential expression. Only genes with FDR<0.05 and an absolute value of logFC > 0.5 were kept for downstream analysis. Heatmaps were generated using ComplexHeatmap (v2.15) (Gu, Eils, and Schlesner 2016).

For performing differential expression of monocyte-derived cells, we used MAST (v1.3) (Finak et al. 2015) as this category had many small populations that were filtered out by edgeR. For microglia, we performed both pseudobulk (as above) and MAST (v1.3) to confirm the validity of the identified clusters.

### Gene Ontology analysis

We used clusterprofiler (version 3.14.3) (Yu et al. 2012), specifically the enrichGO function for over representation analysis, set for biological process (BP) or All ontologies.

### Gene Set Enrichment analysis

The GSEA analysis was run using the fgsea package version 1.12.0 (Korotkevich et al. 2016). Differential expression results for each oligodendrocyte or microglia cluster were ordered by log (fold change). The pathways for each glial subtype were derived from positively differentially expressed genes identified in an oligodendrocyte (Pandey et al. 2022) and microglia metanalysis (Gazestani et al. 2023). Data was plotted using the ComplexHeatmap R package.

### Interferon module score and thresholding

For determining the module score of Interferon response gene signature, we used the murine Type I Interferon gene list from MsigDB. Similar results were observed when a Type II gene list, also from MsigDB, was used (data not shown). This is likely due to the overlap of genes induced by Type I and II Interferon pathways (Liu et al. 2012).

To threshold the Type I Interferon response module score, we considered all cells in the dataset together. First, we removed cells with a negative module score, then calculated the value of the score mean plus two standard deviations. We considered cells above this value (0.348) to be “Interferon-high”. For calculating the differences between Interferon-high cells and all other cells, we performed differential expression using MAST (version 1.3).

### Slide-seqv2: Alignment and Spatial Analysis

Alignment of Slide-seqv2 data was performed using slideseq-tools (https://github.com/MacoskoLab/slideseq-tools). Resulting gene expression matrices and array locations were used to create a Seurat object. Arrays were examined for density of beads and 3 pucks were excluded from the analysis large missing areas in the bead array.

As the bead array covered a large amount of the brain (eg. cortex and hippocampus) we next segmented out the remyelinating lesion or white matter (for controls). For each puck we binarized the expression of *C1qa* and *Mbp*, using this as a map to identify remyelinating white matter (Fig. 1f for example). Segmentation was performed iteratively, first we used the CellSelector function in Seurat, followed by a custom Shiny (v1.8.1) application that enabled a refined selection of the remaining lesion beads (see Fig. 1g and Extended Data Fig. 1f-g). All subsequent analyses were carried out on these cropped arrays.

All fully cropped beads were merged into the same Seurat object and underwent a standard processing pipeline (normalization using NormalizeData function) to generate the final unbiased clustering based on gene expression only (Louvain). We identified 19 clusters from this spatial dataset based on gene expression alone. We performed a differential expression on these clusters using the “MAST” package (v. 1.3). To annotate these clusters we first identified markers of each glial immune cell type by performing a cell type-level differentially expression analysis (MAST, v1.3) and took the top 50 genes per cell type to make “cell type pathways”. Then we used these gene lists in two ways: 1) We performed a gene set enrichment analysis using the fgsea package (version 1.12.0) on the spatial differential expression using the cell type pathways. 2) We applied a module score (using AddModuleScore function from the Seurat package).

To identify the core of the lesion, we calculated a spline along the *C1qa* positive beads using the smooth.spline function (R package stats v4.4) (ref) with df=5. We then calculated the distance for each bead in the lesion from this spline and calculated the average per cluster and timepoint.

### Imaging Analysis

For all histological analyses, lesions were manually segmented based on DAPI hypercellularity in FIJI (v.29.1/1.53t). In order to account for Interferon glia adjacent to the lesion, an additional region-of-interest (ROI) was created for analysis of *Cxcl10* cell density by enlarging the lesion ROI by 50 μm in all directions. Cells within this larger ROI, but outside of the lesion itself were counted as extralesion cells (as opposed to intralesion cells).

For quantification of STAT1 and *Ifit2* signal, mean intensity values in the lesion for each marker were recorded. For quantification of CD3, CD4, CD8, KLRB1C, *C1q, Aldh1l1,* and *Cxcl10* cell density, positive cells were manually counted in Qupath (v.0.5.0 for Mac x64). The vasculature marker CD31 was used with CD3 and KLRB1C stains to determine the proportion of T and NK cells that were truly parenchymal. For quantification *of Pdgfra, Plp1*, and *Aspa* cell density, Qupath was used to segment cells based on DAPI signal, then an object classifier was trained to detect OPCs (*Pdgfra*+/*Plp1–/Aspa–*), premyelinating OLs (*Plp1+/Aspa–*), and myelinating OLs (*Plp1+/Aspa+*). Mean intensity or cell counts per lesion were exported into R (v.4.3.1) for further processing, statistical analysis, and graphing.

Cell counts were divided by lesion area to calculate cell density. Final cell density and mean intensity values per animal were determined by taking an average of two lesion sections, approximately 130 microns apart.

Student’s two-sided t-test was used to determine significance. All plots were made using the ggplot2 (v.3.4.4) package.

### Single-cell RNAseq of iMGs

Cellranger (version 6, 10X Genomics) was used for demultiplexing, barcode preprocessing, generation of fastq files, alignment and the counting of unique molecular identifiers (UMI). We applied Cellranger’s aggr function to all samples to merge and normalize by sequencing depth. We then applied CellBender to remove ambient RNA and technical artifacts due to barcode swapping (Fleming et al. 2023).

Gene expression matrices were analyzed using Seurat (v5) (Hao et al. 2024). We filtered only high quality cells based on the following metrics: Number of genes > 750 and < 6000. UMIs <1500 and>25,000 and percentage mitochondria < 20%, as per our prior work (ref). After this filtration, we identified 55,724 cells. Cells were normalized using the NormalizeData function in Seurat with the LogNormalize option and a 10,000 scale factor. Cells were scaled and clusters were identified based on gene expression using Louvain method.

To identify differentially expressed genes by stimulation (PBS or human IFN-β), we used “edgeR” (v4.2). Raw counts were summed by sample (ie. differentiation) and stimulation using the Seurat2PB function in edgeR. Sample clusters with a library size < 5e4 were filtered out as were lowly expressed genes (using filterByExpr function in edgeR). We performed a TMM normalization, dispersion estimation and differential expression. Only genes with FDR<0.05 and an absolute value of logFC > 0.5 were kept for downstream analysis.

### Bulk RNA seq analysis

FastQC (v.0.12.1) and MultiQC (v.1.23) were to generate initial quality control (QC) reports. STAR (v.2.7.10b) was used to build the reference index (based on GRCh38) and align reads, then the quality of the generated BAM files was assessed with Samtools (v.1.6) quickcheck (all samples had greater than 94% alignment). The count matrix was generated using featureCounts (v.2.0.6).

R (v.4.3.1) was used for subsequent analyses. Differential expression analysis was performed with limma (v.3.56.2), Ensembl gene IDs were mapped to gene symbols using annotables (v.0.2.0), volcano plots and heatmaps were generated with ggplot2 (v.3.4.4), gene ontology (GO) enrichment analysis was performed with clusterProfiler (v.4.8.3) and GO barplots were created with enrichplot (v1.20.3).

---

**Extended Data Figure 1:**
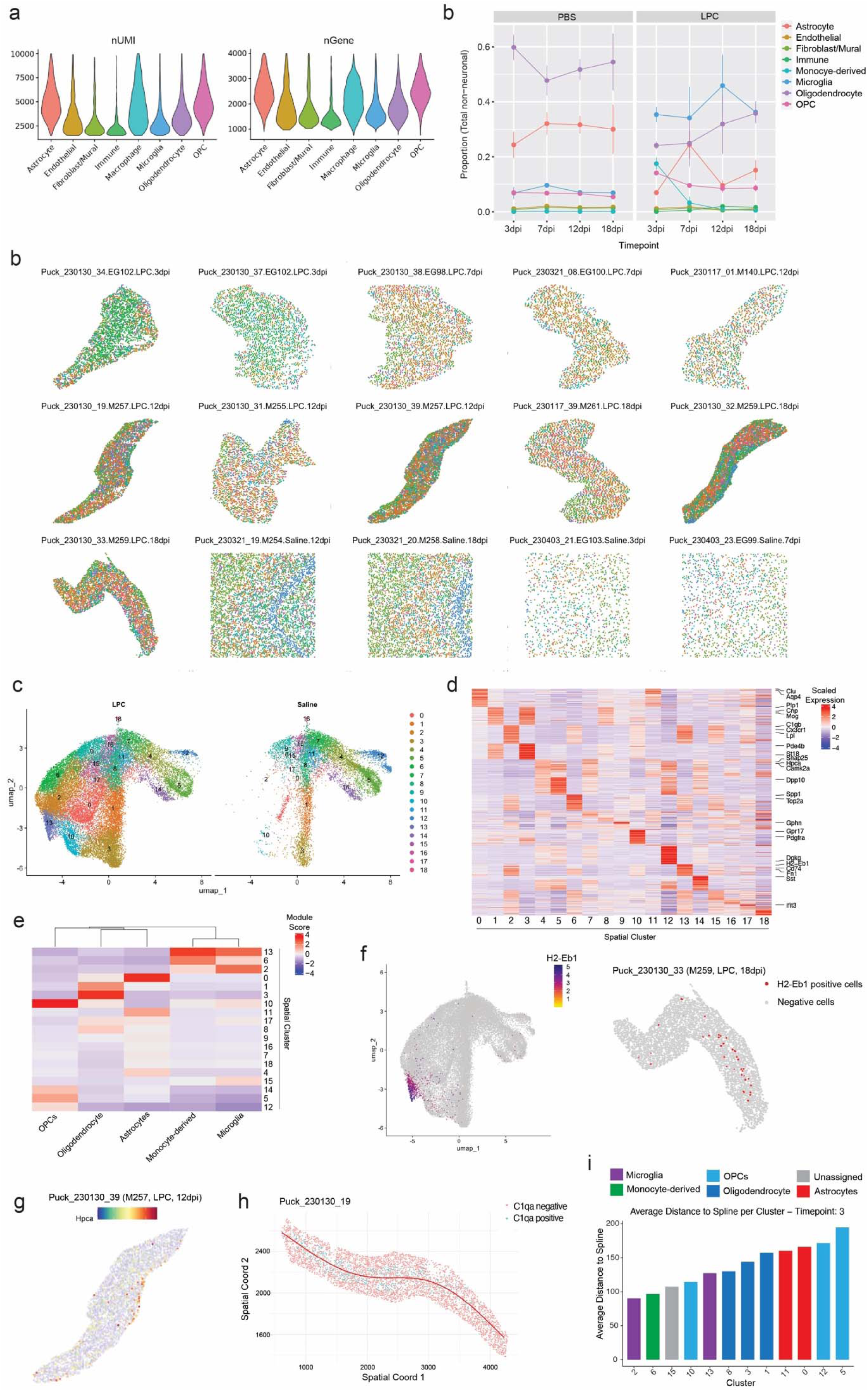
a) Quality control metrics for the full remyelination dataset (excluding neurons) plotted per cell type: nUMI= number of unique molecular identifiers, nGene is number of unique genes expressed. b) All Slide-seqv2 bead arrays (“pucks”) with beads plotted per cluster c) UMAP of all Slide-seqv2 beads plotted by gene expression clustering (“spatial cluster”) and separated by condition. d) Heatmap of top differentially expressed genes per spatial cluster. e) Heatmap of cell type module scores per spatial cluster. f) (Left) UMAP of all Slide-seqv2 beads expressing *H2-Eb1* (Right) Segmented Slide-seqv2 beads from an 18dpi LPC-injected mouse showing *H2-Eb1* expression. g) Slide-seqv2 array showing neuronal gene expression of *Hpca* in the hippocampus. h) Identification of lesion core by tracing a spline along C1qa positive beads. i) Average distance to spline per spatial cluster for 3dpi.

**Extended Data Figure 2:**
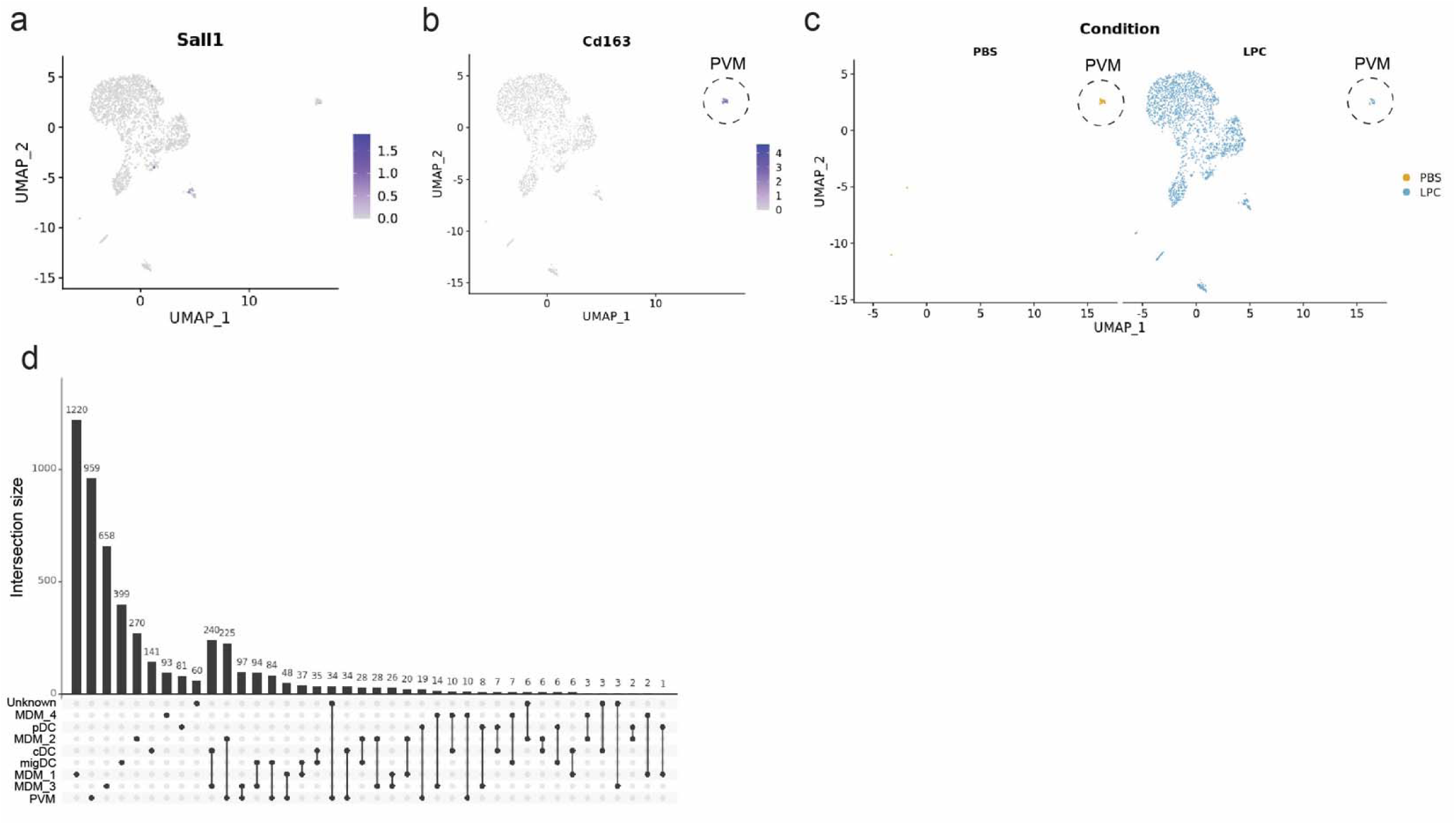
a) UMAP of monocyte-derived cells showing lack of *Sall1* expression. b) UMAP of monocyte-derived cells showing *Cd163* expression with perivascular macrophages (PVM) outlined. c) UMAP of monocyte-derived cells by condition, PBS saline or LPC (remyelination). d) Upset plot showing unique differentially expressed genes per monocyte-derived cluster.

**Extended Data Figure 3:**
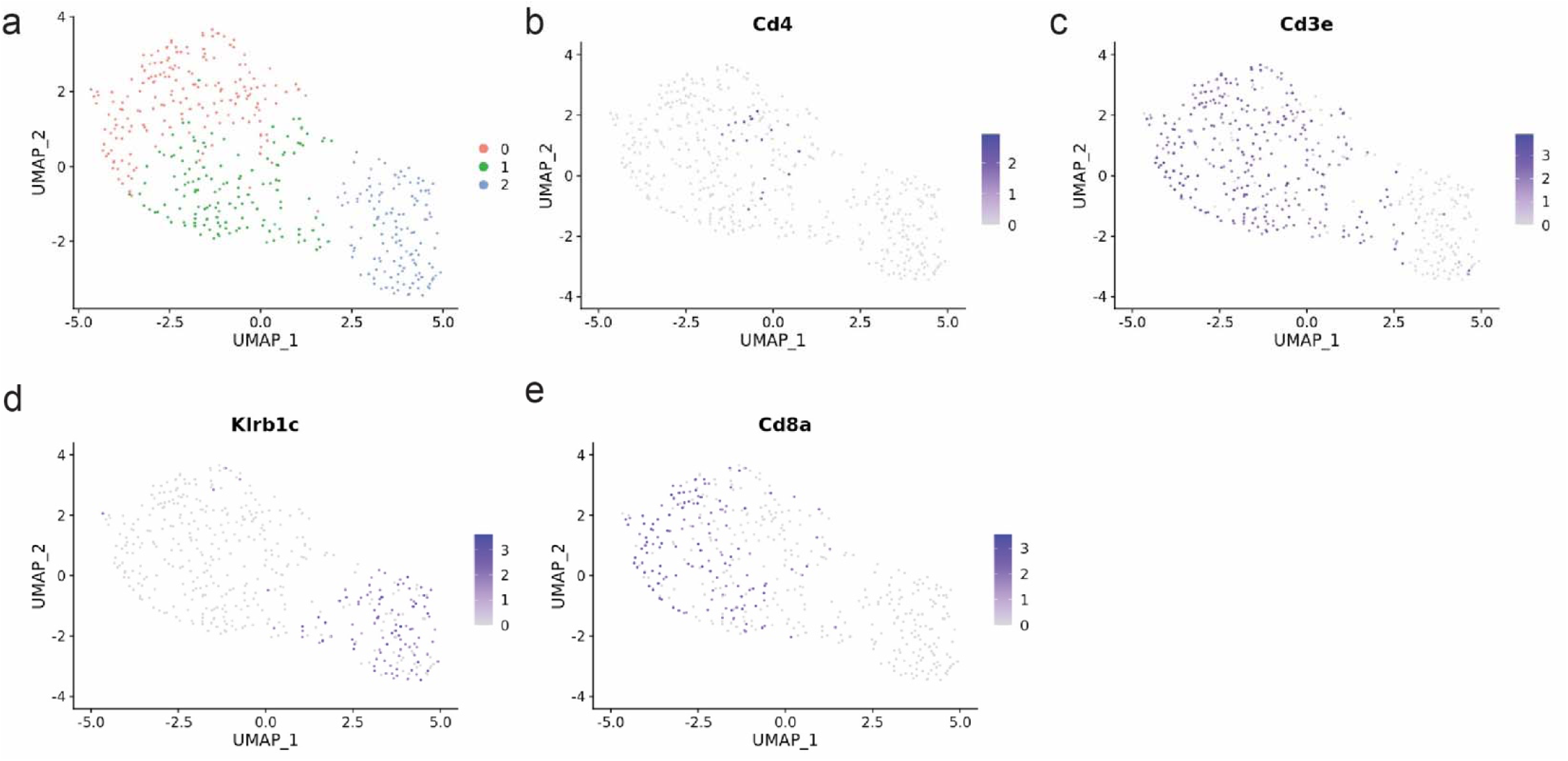
a) UMAP of lymphocytes plotted by cluster. b-e) UMAP of lymphocytes showing expression of *Cd4*, *Cd3e*, *Klrb1c* and *Cd8a*.

**Extended Data Figure 4:**
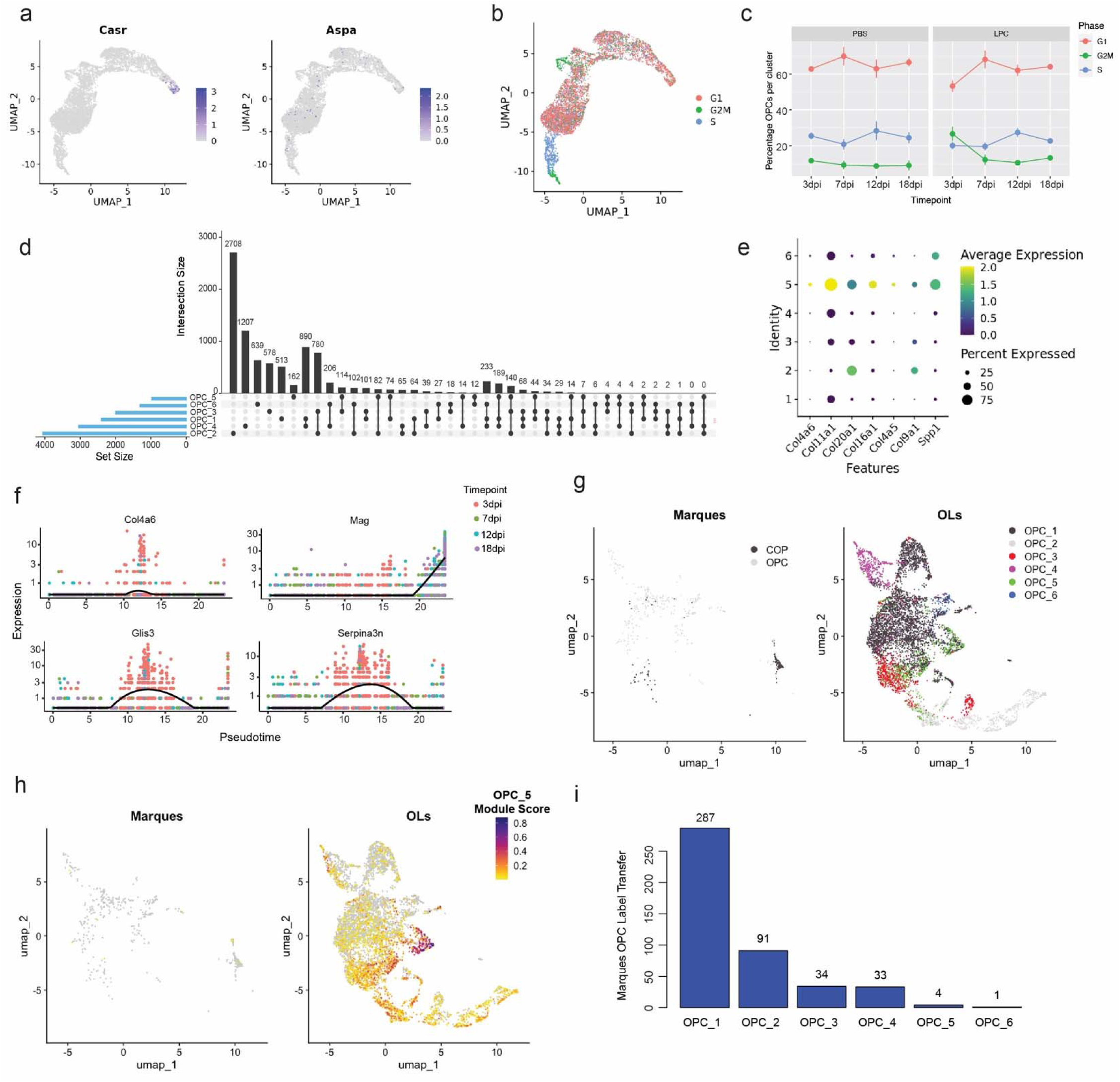
a) UMAP of OPCs and Pre-OLs showing low expression of *CasR* and *Aspa*. b) UMAP of OPCs and Pre-OLs plotted by cell proliferation classification. c) Proportion of OPCs and Pre-OLs plotted by condition and timepoint. d) Upset plot showing unique gene expression per OPC and Pre-OL clusters. e) Dotplot of extracellular matrix genes plotted per OPC and Pre-OL cluster. f) Pseudotime of LPC-injected OPCs and Pre-OLs plotted by timepoint. g) LIGER dataset integration of OPCs and differentiation-committed oligodendrocyte precursors (COPs) from the Marques dataset^39^ with OPCs and Pre-OLs in our snRNAseq dataset. UMAP split by data source. H) UMAP of dataset integration with a module score for genes enriched in OPC_5, split by data source. i) Label transfer of cells from the Marques dataset^39^ (OPCs and COPs only). Only 4 cells are assigned to cluster OPC_5.

**Extended Data Figure 5:**
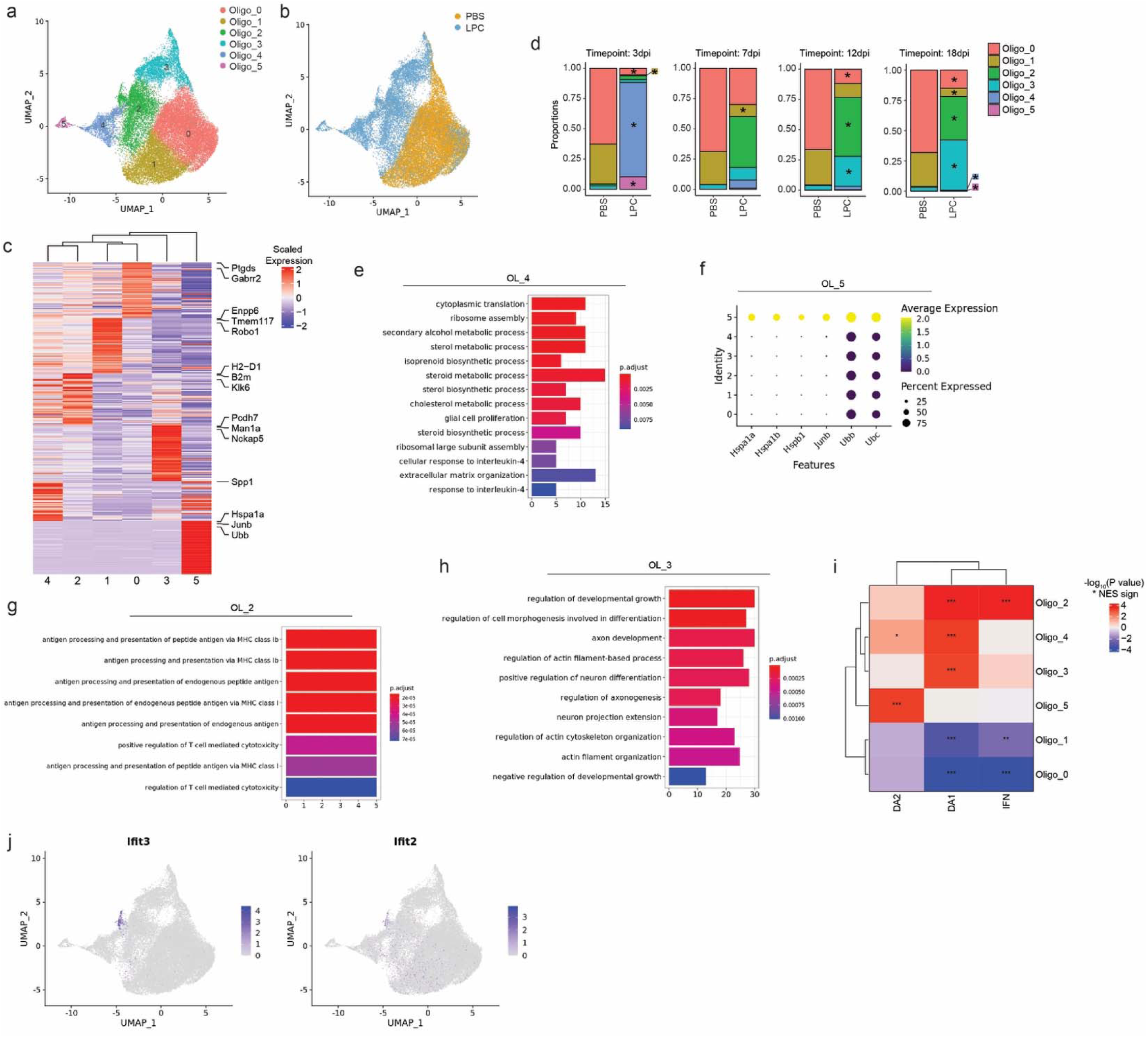
a) UMAP of mature oligodendrocytes labelled by cluster. b) UMAP of mature oligodendrocytes labelled by condition. c) Heatmap of differential gene expression per mature oligodendrocyte cluster. d) Proportion of mature oligodendrocyte clusters per timepoint and condition. * = p-value<0.05 and denotes statistical significance determined by a differential abundance test. e) Gene ontology for genes enriched in OL_4. f) Expression of heat shock and ubiquitin genes expressed per mature oligodendrocyte cluster. g) Gene ontology for genes enriched in OL_2. h) Gene ontology for genes enriched in OL_3. i) Heatmap of the relative GSEA enrichment significance of positively enriched marker genes from disease-associated oligodendrocyte signatures identified in a large-scale meta-analysis^18^. * = p-value<0.05, ** = p-value<0.01 and *** = p-value<0.001. j) UMAP of Interferon responsive gene expression *Ifit3* and *Ifit2* in mature oligodendrocytes.

**Extended Data Figure 6:**
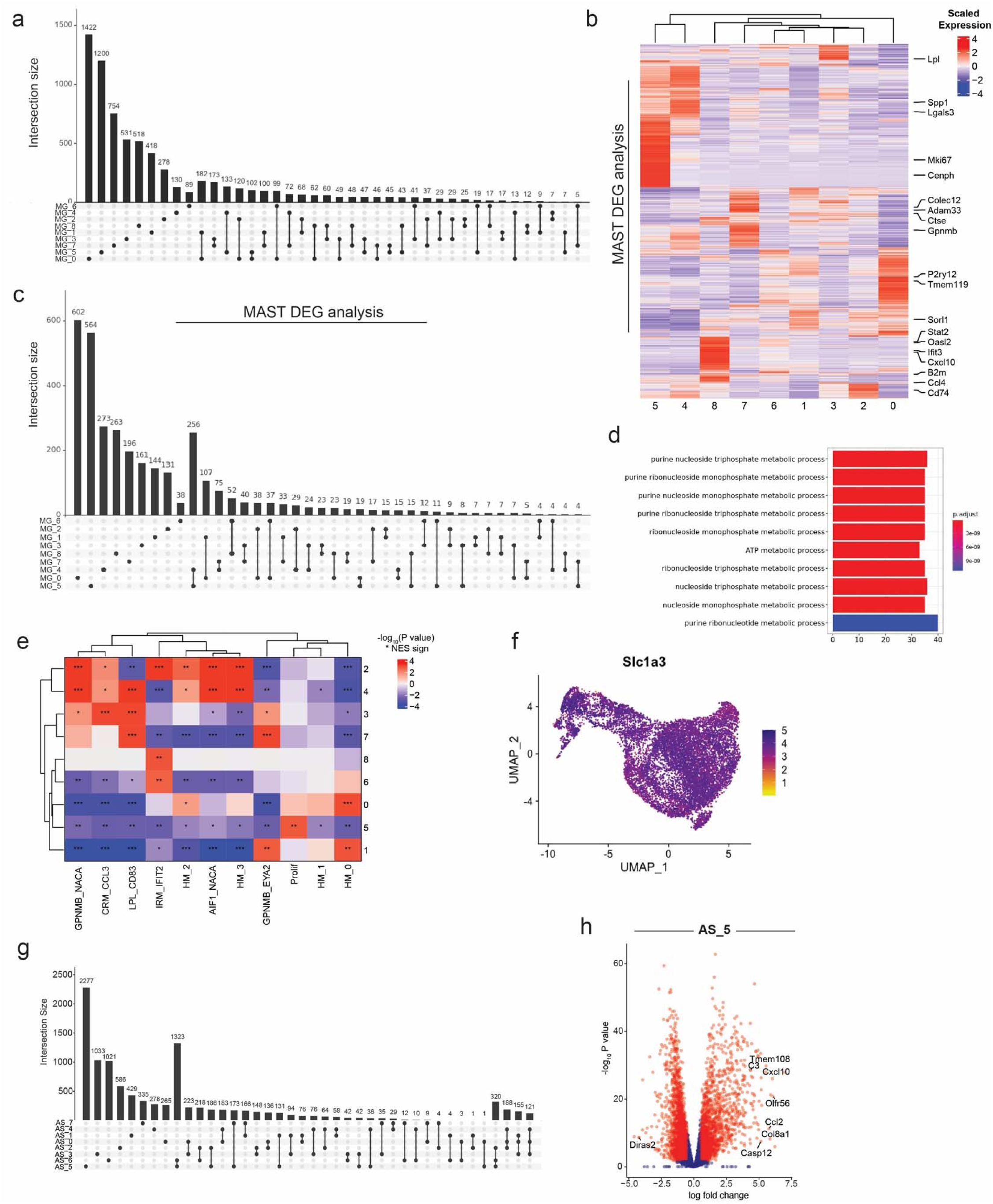
a) Upset plot showing unique differentially expressed genes per microglial cluster (calculated using pseudobulk approach). b) Heatmap of differential gene expression per microglial cluster (determined using MAST^40^). c) Upset plot showing unique differentially expressed genes per microglial cluster (determined using MAST^40^). d) Gene ontology analysis of enriched genes in MG_4 demonstrating enrichment of purine metabolism. e) Heatmap of the relative GSEA enrichment significance of positively enriched marker genes from microglial state signatures identified in a large-scale meta-analysis^42^. * = p-value<0.05, ** = p-value<0.01 and *** = p-value<0.001. f) Expression of *Slc1a3* (*Glast*) in astrocytes. g) Upset plot of unique differentially expressed genes per cluster. h) Volcano plot of remyelination-specific astrocyte cluster AS_5, showing differentially expressed genes relative to other clusters.

**Extended Data Figure 7:**
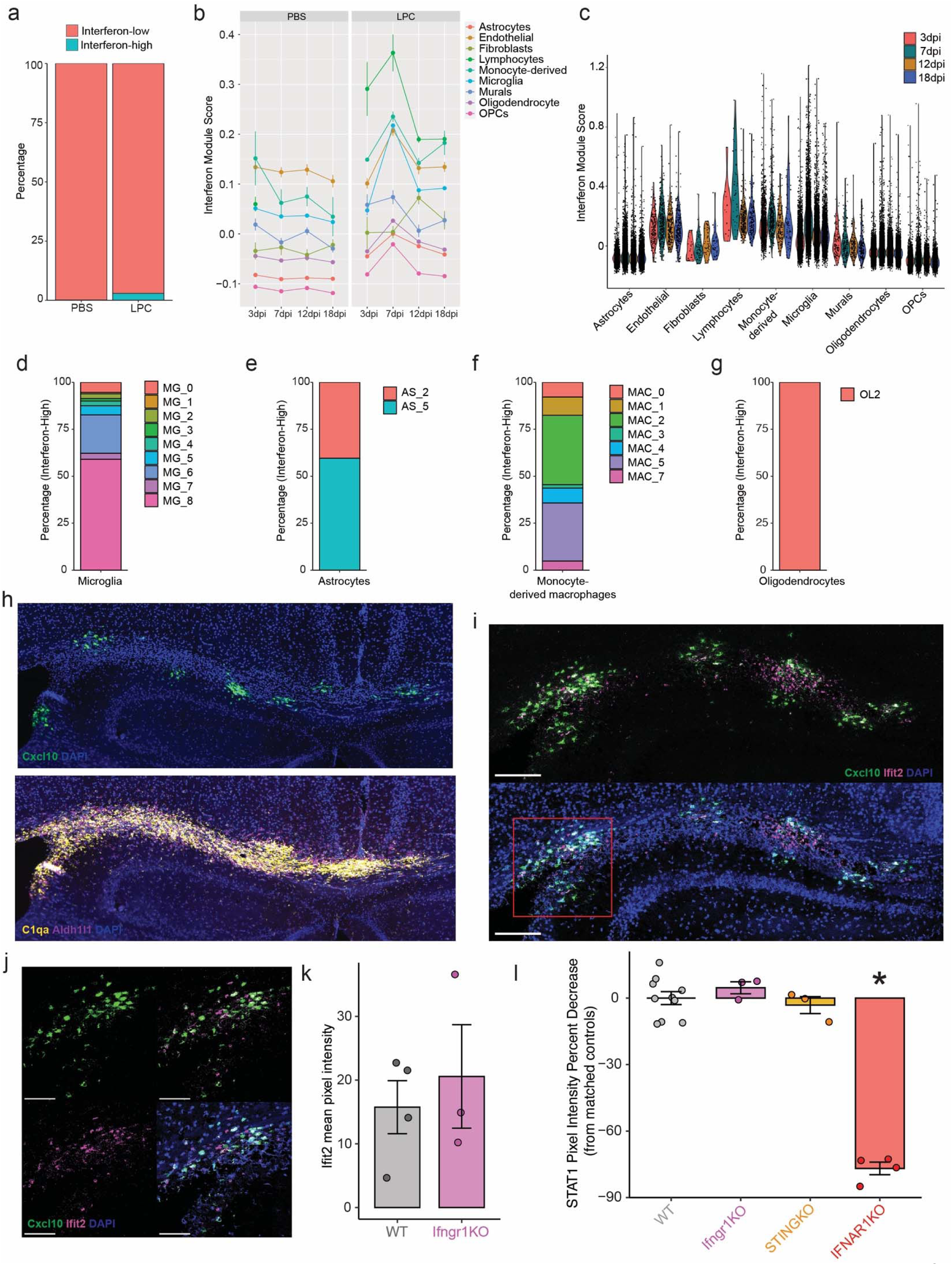
a) Percentage of interferon-low and interferon-high cells split by condition. b) Interferon module score plotted per cell type and timepoint and split by condition. c) Interferon module score in LPC-injected mice plotted per cell type and timepoint. Each point is a cell. d) Proportion of Interferon-high microglia per microglial cluster. e) Proportion of Interferon-high astrocytes per astrocyte cluster. f) Proportion of Interferon-high monocyte-derived cells per cluster. g) Proportion of Interferon-high oligodendrocytes per mature oligodendrocyte cluster. h) Representative *in situ* hybridization for *Cxcl10* mRNA at 7dpi after LPC injection. (Top) Expression of Cxcl10. (Bottom) Expression of microglia and astrocyte cell type markers (*C1qa* and *Aldh1l1* respectively). i) Representative *in situ* hybridization for *Cxcl10* and *Ifit2* mRNA at 7dpi after LPC injection, without (Top) and with (Bottom) DAPI. j) Zoom of inset in (i) showing *in situ* hybridization for *Cxcl10* and *Ifit2* mRNA. k) Quantification of mean *Ifit2* intensity in wild-type and *Ifngr1* knockout mice at 7dpi. l) Quantification of STAT1 immunohistochemistry in the lesions of wild-type (WT), *Ifngr1*, *Sting1* and *Ifnar1* knockout mice at 7dpi. *= p-value<0.05.

**Extended Data Figure 8:**
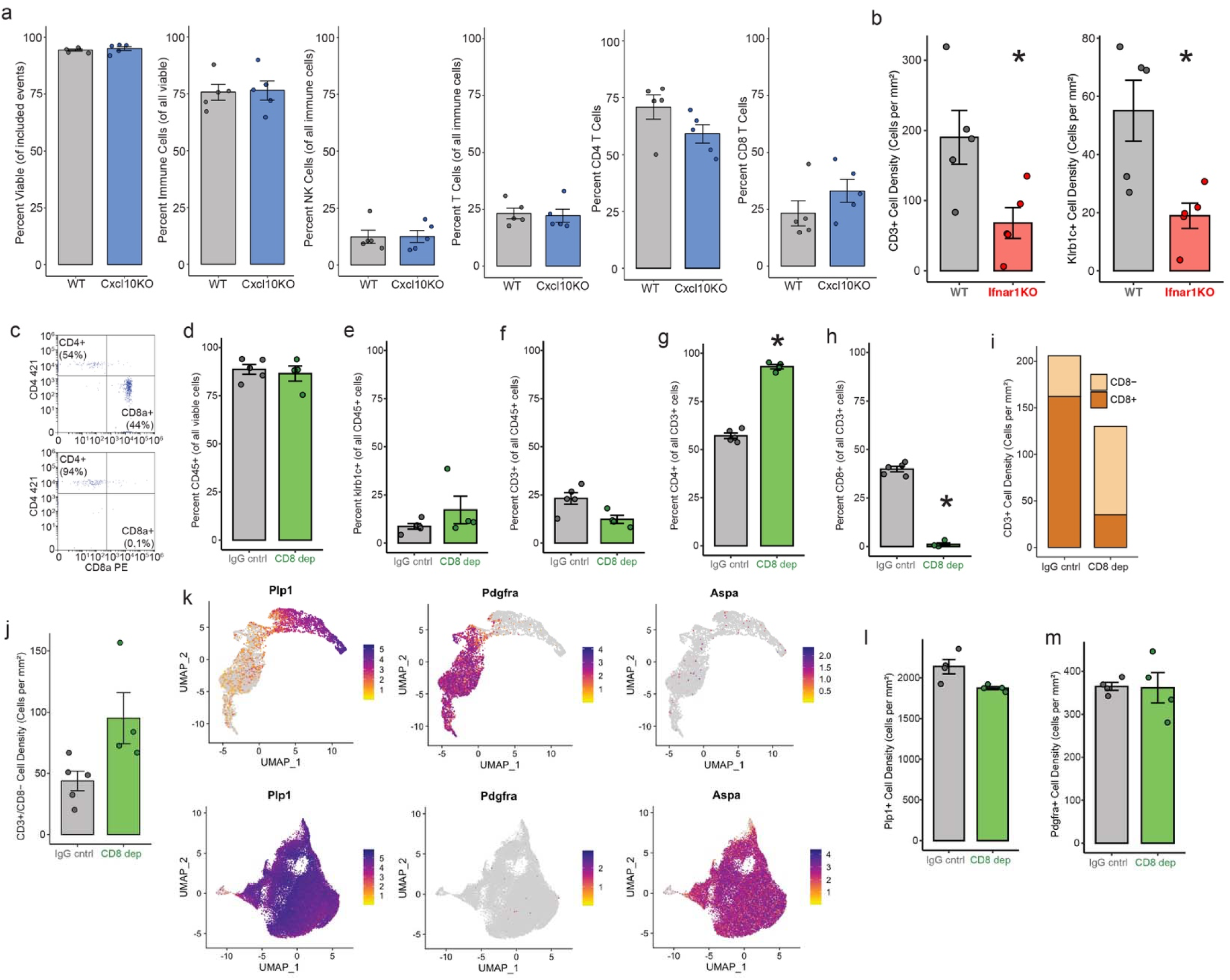
a) Percentage of viable, CD45^+^ immune cells, KLRB1C^+^ NK cells, CD3^+^ T-cells, CD3^+^ CD4^+^ T-cells and CD3^+^ CD8A^+^ T-cells, determined by flow cytometry, from the blood of wild type and Cxcl10 knock out mice at 15dpi after LPC injection. b) Density of T-(left) and NK cells (right) in remyelinating lesions determined by immunofluorescence at 15dpi in wild-type and *Ifnar1* knockout mice. c) Representative flow plot showing CD3^+^ cell expression of CD4^+^ and CD8A^+^ for IgG control (Top) and CD8 depletion (Bottom). d-h) Percentage of blood (d) CD45^+^ immune cells, (e) KLRB1C^+^ NK cells, (f) CD3^+^ T-cells, (g) CD3^+^ CD4^+^ T-cells and (h) CD3^+^ CD8A^+^ T-cells determined by flow cytometry on blood from LPC-injected animals at 18dpi in IgG control or CD8 antibody depleted animals. i) Density of T-cells in remyelinating lesions determined by immunofluorescence at 18dpi in IgG control or CD8 antibody depleted animals. j) Density of CD3^+^ CD8^-^ T cells in remyelinating lesions at 18dpi in IgG control or CD8 antibody depleted animals. k) UMAPs of OPCs and Pre-OLs (Top) and mature oligodendrocytes (Bottom) showing expression of key marker genes using in the *in situ* hybridisation panel: *Plp1*, *Pdgfra* and *Aspa*. l-m) Density of *Plp1*^+^ mRNA (l) or *Pdgfra*^+^ (m) positive cells in remyelinating lesions at 18dpi in IgG control or CD8 antibody depleted animals. *= p-value<0.05.

**Extended Data Figure 9:**
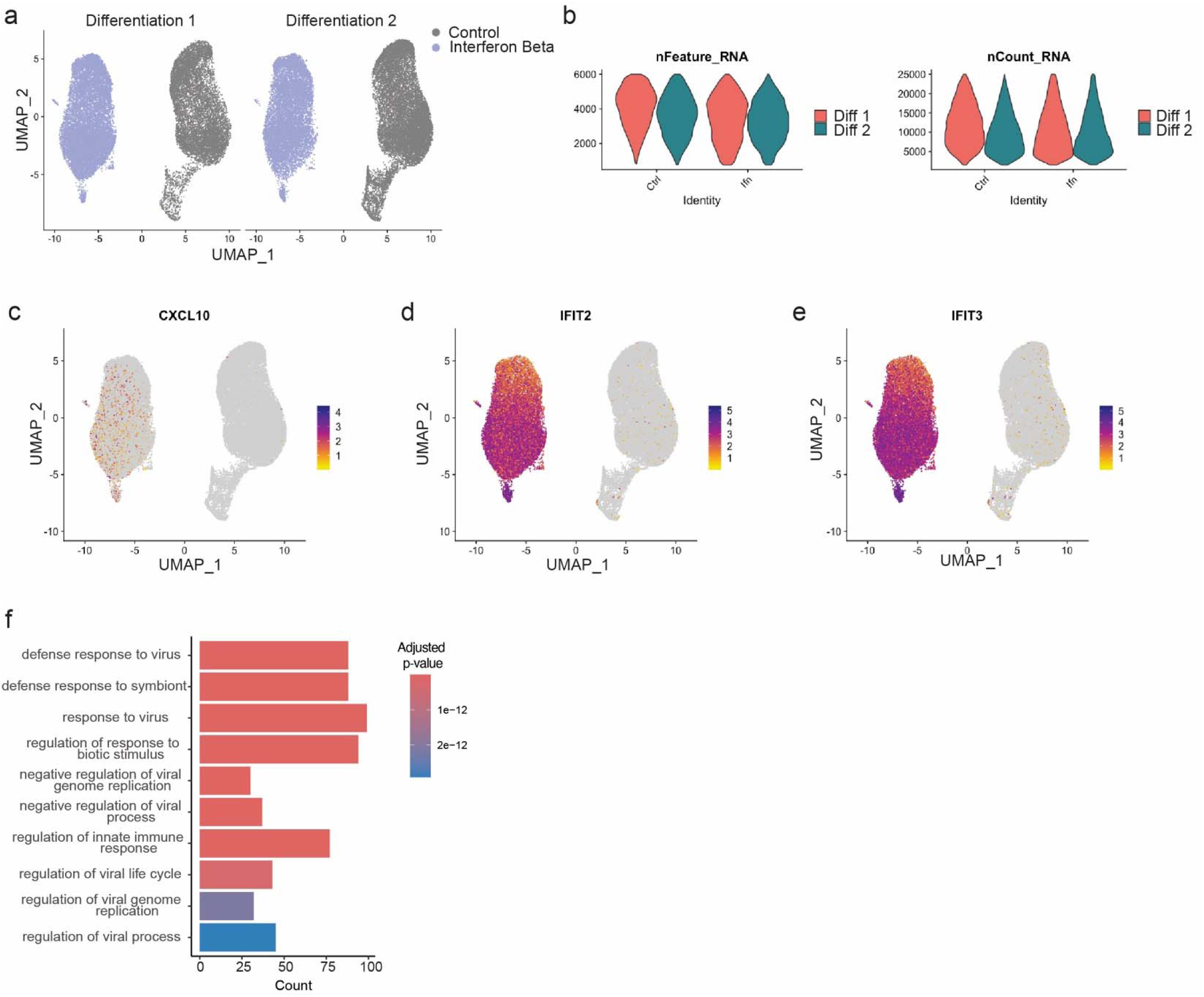
a) UMAP of iMG single-cell RNA sequencing plotted by exposure to saline vehicle or 25ng/ml human Interferon-β, separated by differentiation. b) Single cell quality control metrics per condition and differentiation. nFeature_RNA is the number of unique genes detected in each cell. nCount_RNA is the total number of UMI (unique molecular identifier) counts per cell. c-e) UMAP of *CXCL10*, *IFIT2* and *IFIT3* mRNA expression in iMGs. f) Gene Set Enrichment Analysis of differentially expressed genes enriched in iAstros exposed to 25ng/ml human Interferon-β, compared to vehicle control.

